# Tensile expansion microscopy applies mechanical force to super-resolve fixed and image live cellular samples

**DOI:** 10.64898/2026.02.20.707066

**Authors:** Vignesh Venkataramani, Danielle R. Latham, Ramita Arampongpun, Marouen Zammali, Tejasvin Shrikanth, Anshuman Mohapatra, Jason A. Guerrero, Roberto C. Andresen Eguiluz, Divita Mathur, Laura M. Sanchez, Lydia Kisley

## Abstract

Understanding biophysical phenomena requires techniques that access biologically relevant spatial and temporal scales. Expansion Microscopy (ExM) is a sample preparation approach which achieves super-resolution spatial scales by leveraging osmotic forces in a swellable hydrogel to physically separate structures to distances larger than the diffraction limit of light. Yet, in traditional osmotic ExM only pre- and post-expanded samples can be imaged. Further, fragmentation, hydrogel deformation, and signal loss are common while requiring samples to be chemically fixed. Therefore, there is little control of the expansion, reproducibility can be challenging, and dynamics of biological samples at applicable temporal scales cannot be observed. Here, we develop Tensile Expansion Microscopy (TExM) to mechanically expand fixed and, notably, living cellular samples. Highly-stretchable and tough double network alginate- Ca^2+^/polyacrylamide hydrogels are expanded by tensile forces applied using an electromechanical iris expansion device during continuous imaging on a fluorescence microscope. We incorporate two-photon polymerized microscale fluorescent fiducial markers to track samples and distortion during expansion. The hydrogels controllably and repeatedly expand up to 3.3× with distortions less than 12 *µm* across 1.3 *mm*^2^. TExM is first applied to fixed NIH 3T3 fibroblast cells with immunohistochemistry-stained microtubules, achieving super-resolutions of 100 nm. Then, TExM is demonstrated with living HeLa cells with internal fluorescent reporters showing increased cell size and cell-to-cell separation under 3.2× linear expansion. Overall, TExM allows for continuous, stepwise, and precise temporal modulation of lateral substrate strain, enabling real time monitoring of dynamics of both fixed and viable live cell processes at higher spatial resolutions. TExM can further investigate broad biophysical questions due to its compatibility with other analytical imaging methods that are sensitive to water or fixatives used in traditional osmotic ExM.

## Introduction

Studying biological systems at their native spatial and temporal scales is essential to gain insight into relevant biophysical phenomena. Expansion Microscopy (ExM) has brought innovation at the subcellular to tissue spatial scales, using readily available and inexpensive reagents to enable super-resolution imaging with simple optical setups.^1^ In ExM osmotic forces drive (ideally) isotropic 4-20× expansion of cells and tissues through the uptake of water into hydrogels that are conjugated to the biological sample. ExM is highly adaptable and has enabled the investigation of Alzheimer’s disease,^2^ high-throughput analysis of neuronal circuits and arrangement of pre- and post-synaptic proteins in the brain,^3^ scalable spatial transcriptomics,^4–6^ and metabolic inhomogeneities in tumor tissues through spatial multi-omics profiling.^7^ Although ExM is a versatile and powerful structural imaging modality, control and measurement of the degree and isotropy of expansion is challenging, and the ability to observe the dynamics of the expansion or the temporal evolution of biological samples remains limited.

High structural resolution with isotropic expansion in osmotic ExM is possible only through a combination of the physicochemical properties of the hydrogel composition and sample digestion homogenization. Osmotic ExM methods require chemical fixation of the sample and digestion of the cell wall and protein structure. High quality reagents are required in specific ratios to obtain larger expansion factors consistently.^1,8^ Uniform sample digestion is critical to obtaining isotropic expansion.^8–11^ Digestion breaks the fundamental covalent bonds of the sample, chemically homogenizing the sample. However, the digestion step is hard to optimize due to variability within biological samples,^12^ stability of macro-molecules and fluorophores of interest, ^13^ and the polymer system used for expansion.^14,15^ This variability limits the reproducibility of the technique, as the unique biological features of each sample dictates specific digestion requirements that may vary between experiments. The quantification of isotropy is often done using intrinsic reference structures within the biological sample itself. Using intrinsic reference structures leads to a lack of robust quantification in regions where reference structures are not present. Recent studies have shown the benefit of using external reference structures, or fiducial markers, to obtain local expansion factors and correct for local distortion,^11,16^ but still suffer from loss of signal from digestion and volumetric dilution upon expansion. This shows a need for a hydrogel system that can be reproducibly and controllably expanded with robust extrinsic markers to track expansion isotropy.

Osmotic forces drive the expansion of the hydrogel network in ExM, limiting the controllability, repeatability, and reversibility of the expansion.^17,18^ After fixation and digestion, expansion is performed by soaking the sample in water or a buffer prior to immobilizing the sample on a microscope-compatible coverslip for imaging. Therefore, only the pre-expansion and post-expansion data can be collected; the expansion cannot be easily monitored or stopped at a given expansion factor. Thus, the dynamic osmotic expansion step is not compatible with microscopy. Any potential anisotropic displacement of the fragmented structural features during the expansion process cannot be observed.

The structural organization of macromolecules or cells is only observed in osmotic ExM while the biological sample is in a static state.^17,18^ A key step in osmotic ExM and its variants^10,14,19,20^ is the required fixation of the sample to preserve the sample configuration, effectively taking a snapshot in time.^9,21^ Fixing crosslinks biomolecules of interest into place making the study of any cellular or intermolecular dynamics impossible to achieve. Fixation protocols can also lead to artifacts, disrupting protein-protein interactions.^22^ Notably, fixation makes ExM methods fundamentally incompatible with live cell imaging. In contrast, currently available methods for super-resolution, dynamic imaging of live cells or the cellular environment such as PALM, STED, MINFLUX or fcsSOFI,^23–26^ in situ require more expensive equipment and can be technically challenging in either sample preparation or computational image post-processing.

Here, we report Tensile Expansion Microscopy (TExM) which employs tensile forces, applied through a custom iris-based expansion device to equi-multiaxially expand a mechanically extensible and tough hydrogel system incorporated with microscopic fluorescent fiducial markers alongside biological samples. The application of tensile force through the iris expansion device onto the double network hydrogel is controlled using a stepper motor and micro-controller chip, making the expansion precise, repeatable and reversible. By incorporating microscale fiducial markers in the sample, we are able to continuously monitor the same region over the expansion while being able to quantify anisotropy with higher precision without requiring fluorescence signal amplification. Electronically controlling the iris-based mechanical expansion device to expand polymeric double network hydrogels,^27^ we show minimal distortion in expansion over areas in the order of 10, 000 *µm*^2^. Using tensile forces instead of osmotic forces, we control and track the degree of expansion along with the rate of expansion in real time. By continuously monitoring the expansion of fixed NIH-3T3 cells,^28^ we show improvement of spatial resolution as we increase expansion to achieve super-resolution. Further, our method using tensile forces to expand allows the study of live cell systems. We show an increase in cellular size of HeLa cells during expansion using real time imaging. We report TExM can achieve an optical resolution of 183 nm on a 20x (0.5 NA) or 62 nm with a 100x (1.49 NA) objective over 3.3× expansion based on the Rayleigh resolution limit 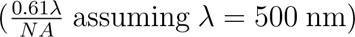. Using a custom expansion device on tough and extensible hydrogel system with fluorescent fiducial markers, TExM enables super-resolution imaging through controlled, repeatable and isotropic expansion of fixed cells while providing a framework for the continuous monitoring of live cell dynamics. TExM is further readily adaptable to a broad range of microscope configurations: upright and inverted, epi- and trans-illumination, with a low-profile design which does not impede the optical path. The iris expansion device assembly uses lightweight thermoplastic and acrylic components, allowing for a high degree of customization while remaining economical and easy to manufacture.

## Results and discussion

### Development of Tensile Expansion Microscopy (TExM)

A highly stretchable, tough, and biologically-compatible alginate-Ca^2+^/polyacrylamide (Alg-Ca^2+^/PAAm) polymeric double network (DN) formulation allows for repeatable and controllable multiaxial expansion (Fig. 1a, i-iv). Adapted from the work of Sun et al.,^27^ we use ionically-crosslinked Alg (Fig. 1a, square inset) and covalently crosslinked PAAm (Fig. 1a, circle inset), which results in a hydrogel that has high stretchability and toughness due to synergy between individual components. During expansion, the Alg-Ca^2+^, a brittle network, acts as a sacrificial component. The relatively weak ionic crosslinks between the Alg chains break and dissipate energy effectively, allowing the hydrogel to absorb significant energy and delay crack propagation. The second ductile PAAm network remains intact and bears the majority of the mechanical load. In its relaxed state, the PAAm network is highly coiled. As the hydrogel expands, these polymer chains are stretched and uncoil. While its strong covalent crosslinks prevent catastrophic fracture and maintain overall structural integrity, the PAAm network undergoes an elasto-plastic deformation. Under equi-multiaxial strain, a considerable amount of these covalent bonds are permanently ruptured, resulting in substantial internal damage that prevents the hydrogel from fully recovering to its original state once the load is removed. The strong covalent crosslinks of the PAAm network prevent the hydrogel from catastrophic fracture, maintaining the overall structural integrity of the material. Further, Alg-Ca^2+^ and PAAm are biocompatible hydrogel substrates commonly used in cell culture.^29,30^ The gelation protocol (Fig. 1a) was modified by tuning the concentrations of the covalent and ionic crosslinkers. This allows modulation of the extensibility, toughness, and stiffness of the hydrogel resulting in hydrogels that can consistently expand 4× multiaxially on the macroscale.

**Figure 1:**
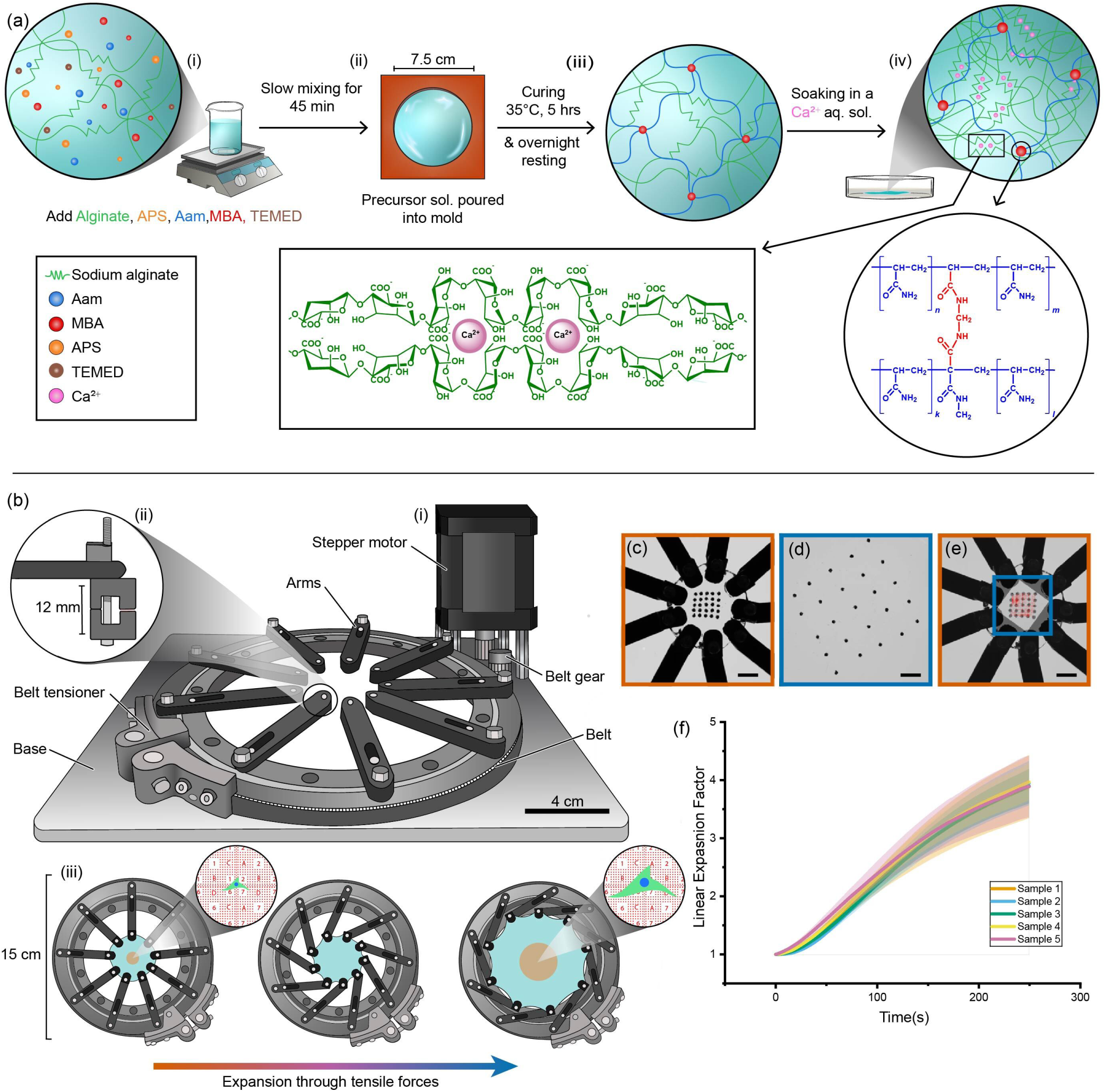
Tensile expansion microscopy (TExM) uses a double network hydrogel and iris based expansion device to achieve up to 4*×* expansion on a macroscopic scale. (a) Schematic of workflow of DN Alg-Ca^2+^ /PAAm hydrogel synthesis compatible with high mechanical strain. (i) A homogeneous precursor solution of Alg, AAm, APS, MBA and TEMED is mixed slowly for 45 minutes. (ii) The solution is poured into a mold and (iii) polymerized by curing at 35*^◦^*C for 5 hours and resting overnight. (iv) The resultant gel is ionically crosslinked by soaking in a CaSO_4_ slurry, producing a gel that is tough enough to handle with at least 4*×* multiaxial expansion. (b) Iris expansion device schematic: (i) A controllable stepper motor rotates nine evenly spaced arms, mounted on a base, through a belt mechanism. (ii) Grippers fit into the end of the arms that secure the hydrogel to the mechanical stretcher. (iii) This mechanism allows hydrogels mounted on the iris to be expanded in a controlled manner. (c) Pre-expansion image and (d) post-expansion image of macroscopic fiducial markers on hydrogel (14 wt.%) loaded on the iris expansion device. Both (c), (d) are cropped to represent the same spacial area. (e) The post-expansion image registered to the pre-expansion image via a similarity transform (representing an isotropic expansion), with a vector deformation field represented by red arrows (see SI method and Figs. S3-5). (c-e) Scale bars represent 10 mm. (f) Tracking and triangulating the fiducial markers over time (see Figs. S4-5) calculates the average linear expansion factor between features (solid lines) and the standard deviation of expansion factors (shaded region). Illustrations in (a, b) attributed to Allison Breuler, reuse copyright CC BY-NC-ND 4.0

In preparation of the hydrogels, the free radical polymerization mechanism is especially sensitive to oxygen.^19,31,32^ We observe near-complete polymerization in about 7 minutes in an argon atmosphere (Fig. S1). For a more accessible and approachable procedure, our formulation accounts for the action of atmospheric oxygen by adjusting the concentrations of the free-radical polymerization initiator and catalyzer (ammonium persulfate (APS) and N,N,N’,N’-tetramethylethylenediamine (TEMED), respectively) (Fig. 1a). This avoids working within an inert glovebox atmosphere, which can be expensive and cumbersome, or semi-permeable glovebags and sample covers, which can lack reproducibility.^19,31^

An iris-design-inspired mechanical expansion device is engineered to apply isotropic multiaxial tensile forces to the hydrogel system in a controlled, reversible, and repeatable manner during optical microscopy. Iris stretchers used for mechanobiology applications^33–35^ are capable of applying physiologically relevant area strain on the order of 100%. To achieve expansion in the order of osmotic ExM, the device was designed to accommodate areal strain in the x–y plane of *ε*_area_ = 1500%, with a large range of motion for the iris arms (Fig. 1b, iii). This corresponds to a 4× linear expansion. Because areal strain can be unintuitive compared to linear expansion factors used in ExM, we report results throughout the manuscript in terms of linear expansion ratios. The iris is built using lightweight and inexpensive materials such as 3D printed polylactic acid plastic and laser cut acrylic. The flat design of the iris can be modified to fit different upright, inverted, epi, or transmission imaging modalities without impeding the optics (Fig. 1b, i-ii). Additional parts for the iris expansion device for hydrogel loading and sample preservation can be found in Fig. S2.

The mechanical iris stretcher is controlled electronically, allowing for precise, repeatable, and fast movement of the iris arms resulting in a desired expansion of the sample. Using custom code to operate an esp32 controller chip and a stepper motor controller mounted on a custom printed circuit board (PCB), we control the movements of the iris arms. During operation, a stepper motor drives a belt, which in turn moves the arms of the iris in a synchronized, precise, and controllable manner. Hydrogel grippers are designed to fit into the end of the stretcher arms and freely rotate during expansion to prevent the application of additional anisotropic strain to the hydrogels while c-clamps are introduced to prevent the grippers from falling off the iris arms (Fig. 1b, ii). The grippers have sandpaper (grit size 60) attached to the surface holding the hydrogel to prevent slipping during expansion. The gripping area needs to be large enough so that the applied pressure does not puncture the hydrogel, while being small enough to maximize expansion. We found that a rounded rectangular area (1 mm x 0.3 mm) works best. The iris arms can be controlled to move further away from each other, increasing mechanical deformation, leading to larger expansion factors (Fig. 1b, iii). The overall range of the arms based on the baseplate size limits the theoretical highest expansion to 4× linear expansion, the motor rotation can expand samples in as low as 0.003× per step, and speed can expand samples on the order of 0.01–5 cm/s. The ability to control and configure the settings of the device offers the versatility required for a broad spectrum of controlled experimental conditions.

We reproducibly and controllably track continuous expansion up to 4× using macroscopic fiducial markers. Here, we use a hydrogel having a total polymer concentration of 14 wt.% (denoted as hydrogel (14 wt.%), see Methods) and AAm/Alg ratio equal to 8.03. The hydrogel is loaded onto the iris expansion device and 2 mm hole punch paper cutouts are placed in a grid pattern on the hydrogel (14 wt.%) and used as fiducial markers. The fiducial markers are held in place by capillary forces between the paper cutouts and the hydrated gel surface. While this interface provides sufficient stability for tracking global expansion, it is possible that micro-scale displacements beyond the threshold of visual detection occur during the expansion process. Such localized movements likely contribute to the baseline error measured in our tracking, though the high reproducibility of our global expansion curves indicates that these effects are minimal. A time lapse video of the hydrogel (14 wt.%) expanding is captured. At any given time, a post-expansion image (Fig. 1d) can be taken and quantitatively compared to the pre-expansion image (Fig. 1c) through an image registration algorithm (Figs. S3-5) to obtain an expansion factor and a deformation field (Fig. 1e, red arrows). The Euclidean distances between neighboring tracked features are calculated and compared to the pre-expansion distances to obtain the average and standard deviation of the linear expansion factor at various expansions. To assess reproducibility, we compared the temporal evolution the expansion of several independent hydrogels (n = 5). The expansion factor for each sample was derived from the mean inter-marker distances calculated via Delaunay triangulation of the fiducial grid (SI Methods, Fig S3). The resulting time-course curves were statistically indistinguishable, confirming a high degree of control over the expansion. Applying the maximum mechanical deformation, within the range of the iris arms, we observe a maximum expansion of 4.1 ± 0.2× with a root mean square error of 220 ± 20 *µ*m. Subsequent experiments focused on the hydrogel formulation with 16.4 wt.% total polymer concentration and AAm/Alg = 8.03, hereafter simply referred to as the “hydrogel”. This change was made to increase hydrogel stiffness and overall cell adhesion.

### Fluorescent two-photon lithography fiducial markers validate Tensile Expansion Microscopy (TExM) at the microscale

Expansion microscopy is performed at the nano to micrometer scales, so we test the expansion factor and local anisotropies at the microscale with nanofabricated fiducial markers. Two-photon lithography features are incorporated into the hydrogel as an independent reference markers to track the local microscopic expansion and distortion of samples. Other works have highlighted the importance of independent reference markers for ExM by quantifying expansion locally in a region where no biological samples are present.^11,16^ IP-Visio, a negative tone photoresist, is a low auto-fluorescence resin designed to be used for fluorescence microscopy. Atto-633 minimizes cross talk to bluer channels during multi-colored imaging of cells. Thus, IP-Visio resin doped with 10 *µ*M Atto-633, is polymerized using a two-photon process in a grid pattern with numbered and lettered identifying features on a flat substrate (Fig. 2a, i,ii, Fig. S6). A hydrogel is cast on top of the substrate and the grid patterns are incorporated into the hydrogel by crosslinking the polymer around the patterns (Fig. 2a, iii). The grid pattern is transferred from the substrate into the hydrogel by carefully peeling the hydrogel (see Methods). Once the hydrogel is ionically crosslinked by soaking in a CaSO_4_ slurry (Fig. 2a, iv), the hydrogel containing the fluorescent fiducial markers is ready for expansion (Fig. 2a, v).

**Figure 2:**
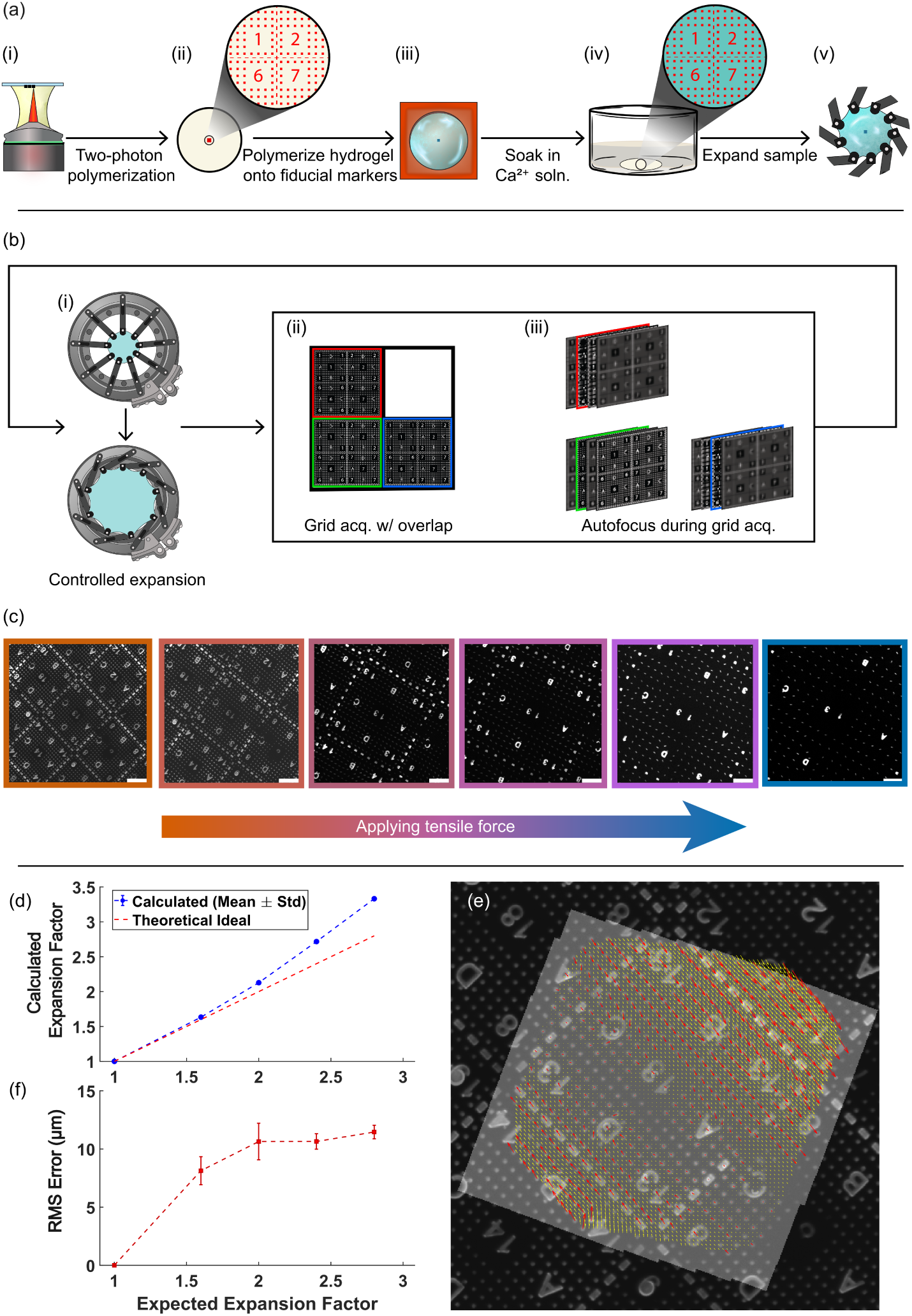
Microscale fluorescent fiducial markers facilitate microscopic imaging of tensile-based expansion. (a) Schematic of workflow to incorporate fluorescent fiducial markers in hydrogels. (i-ii) Microscale fiducial markers are printed on a clean Si wafer through a two-photon lithography process using resin doped with fluorophores. (iii) Hydrogels are polymerized on and around the fluorescent fiducial markers. (iv) The hydrogel is ionically crosslinked in CaSO_4_ slurry to improve toughness resulting in a (v) hydrogel with desirable physical properties for expansion with trackable fluorescent fiducial markers. (b) Flowchart describing the in-focus, MFOV data acquisition during expansion. (i) Iris expansion device is electronically controlled and expanded in 0.5*×* expansion steps. At each step, (ii) multiple images with 80% overlap are acquired in a grid pattern. (iii) At each one of these grid locations, a Python script auto focuses the objective position to acquire an in-focus image. (c) Images with overlap are stitched together to visualize a larger area with fluorescent fiducial markers at every expansion step. Scale bars are 100 *µ*m (d) The measured expansion factor is quantified based on image registration of a post-expansion to a pre-expansion image through a similarity transform at each expansion step compared to the expected expansion factor entered on the iris device electronics. (e) The estimated similarity transform is visualized by overlaying the transformed post-expansion image (gray) on the pre-expansion image (black). Quantified distortion is visualized using red arrows, representing the mismatch between the fluorescent fiducial markers. A vector deformation field is extrapolated and represented by the yellow arrows. (f) The RMS error in expansion is estimated at each expansion step based on the error in the similarity transform obtained. Illustrations in (a,b) attributed to Allison Breuler, reuse copyright CC BY-NC-ND 4.0

Stringent hardware constraints inherent to the fabrication and imaging workflows necessitate the use of specialized substrates capable of maintaining near-atomic flatness and desired optical properties. When imaging, it is beneficial to have biological samples and fiducial markers within the same imaging plane. For example, imaging during expansion with a 20x (0.5 NA) air objective, the depth of field is 1.9 *µ*m, limiting the surface roughness of the hydrogel to *R_a_* ≈ 1.9 *µ*m, with subsequently lower roughnesses required for higher magnification objectives. The hydrogel surface topography is dependent on the flatness of the substrate upon which the hydrogel is set (Fig. S7), and the surface-hydrogel interactions. Additionally, the substrate on which the fiducial marker features are fabricated must have a minimum refractive index difference of Δ*n* = 0.1 with the IP-Visio (*n* = 1.5) for the lithography instrumentation to detect the axial position of the substrate interface.^36,37^ Because the hydrogel’s surface topography mirrors the underlying substrate, we selected silicon (Si) wafers and indium tin oxide (ITO)-coated glass coverslips. Silicon wafers exhibit near-atomic flatness and low warp (*<* 100 *µ*m across 200 mm).^38,39^ Furthermore, they meet the necessary refractive index specifications and are available in diameters exceeding the 30 mm, ensuring compatibility with the iris expansion device (Fig. 1b). 30 mm diameter glass coverslips also work for our purposes with the added benefit of optically transparency and fitting into 6-well plates for cell culture. Cleaned glass coverslips have subnanometer roughness^40^ before being Direct Current (DC) sputter coated with 100 nm of ITO (see Methods) to create a refractive index mismatch for lithography. Alternatively, commercially coated substrates may be used to yield similar results. For the investigation of TExM on the microscale, Si wafers are used.

The physical and photophysical properties of the nanofabricated fiducial markers allow for tracking of unique, identifiable features throughout the expansion process. The Young’s modulus of the IP-Visio resin is multiple orders of magnitude larger than that of the hydrogel systems.^27,37,41^ Therefore, the fiducial markers maintain their size but move away from each other during expansion (Fig. 2c). The fluorescence signal within a single marker is not diluted upon expansion, and we do not observe significant dye diffusion out of the developed resin (Fig. S8). The fluorescence signal does not need amplification after chemical treatment, in contrast to protein-based markers,^11,16^ as a result of the higher stability and quantum yield of the chemical dye. The versitility of these nanofabricated fiducial markers allows for synthesis with different fluorophores at varied concentrations and the use of alternative auto-fluorescent resins (Figs. S9-11). Tracking the fluorescence signal from these markers, we can measure the local expansion factors and anisotropies in the expansion independent of the sample structure.

Coordinated electronic control of the iris stretcher, microscope, and image analysis achieves automatic and in-focus data acquisition during expansion. A single field of view using a 20x air objective to monitor the fiducial markers during expansion is 700 *µ*m x 700 *µ*m. During the tensile expansion process, we gain higher resolution, but are limited to imaging a single, smaller portion of the sample as seen by markers moving away from one another and out of the field of view (Figs. 2c, S10). To quantify the expansion over a larger area, we obtain full field of view images at multiple field of views (MFOV) in a 5 x 5 grid pattern with 80% overlap (Fig. 2b, ii) at each 0.5× expansion step (1×, 1.5×, 2×, …) (see Methods). Due to the hydrogel thinning and heterogeneity in the hydrogel structure, each field of view is located at a slightly different focal plane (Figs. S12-13). We further implemented a custom Python image analysis (see Methods) in the data acquisition loop to keep the microscopic fiducial markers in focus by adjusting the z-position of the microscope objective based on an image sharpness metric (Fig. 2b, iii).

TExM achieves low distortion at the microscale. The acquired MFOV images are stitched to visualize larger areas at each expansion step (Fig. 2b). Pre- and post-expansion images are binarized to identify and correlate the fluorescent fiducial markers (see Methods). Expansion tracking is performed using a MATLAB-based registration function to estimate the similarity transform representing isotropic expansion. For micro-expansion experiments, this transform alone is used to calculate the expansion factor, with statistics derived from repeated estimations (see Methods). For macro-expansion (Fig 1c-f), the expansion factor is determined primarily by a triangulation mesh (described in the previous section), with statistics gathered from the large population of tracked vertices. A similarity transform estimate is primarily used to register pre- and post-expansion images (Fig. S4) but also to confirm that the expansion factor calculated based on triangulation is similar. Using a triangulation based linear interpolation of the mismatch between corresponding pre-expansion and similarity transformed post-expansion fiducial markers, we obtain a vector deformation field (Fig. 2e) corresponding to the non-rigid registration. In both the macro- and micro- scale expansion experiments, sample distortion is quantified via a triangulation-based linear interpolation model applied to the known mismatch between fiducial markers pre-expansion and similarity transformed post-expansion. This allowed us to compare the measured expansion (Fig. 2d) and distortion (Fig. 2f) at an expected expansion programmed into the iris expansion device. We observe a linear trend in the measured expansion factor compared to the expected expansion entered into the iris expansion device electronics, demonstrating the fine control of expansion provided by the iris expansion device. The maximum measured expansion factor based on the similarity transform is above the theoretical ideal (based on the expansion input provided to the iris expansion device electronics) at larger expansions, with a maximum of 3.33× achieved. However, this systematic difference may be within the engineering tolerances of the macroscale motor and 3D printed components of the iris. We also find, at 3.33× expansion the global RMS distortion is 11.5 ± 0.5 *µm* (Fig. 2f). Potential inequalities in the grip of the hydrogels, the iris arms, and larger tolerance gaps in the expansion device may lead to larger variation in distortion across the sample. Overall, both the macroscale and microscale tracking demonstrate that the combination of the DN hydrogel and iris expansion device provides a robust platform for TExM with high reproducibility and minimal local systematic error.

### Super-resolution imaging of fixed cells through Tensile Expansion Microscopy (TExM)

There exist many osmotic ExM protocols for fixed cells and tissues.^1,10,19,21,42^ To equate our TExM approach with osmotic ExM, we apply the iris expansion device to fixed cell samples. Similar to osmotic ExM protocols, our work flow consists of fixing and staining, anchoring, gelation, uplifting and digestion, followed by the expansion (Fig. 3a). Expansion is observed on the microscope with a lower 20-50x magnification air objective to avoid perturbing the hydrogel during the expansion process. At the end of expansion, a drying ring (Fig. S2c) can be used to immobilize and preserve the hydrogel in an expanded state allowing for further imaging with higher resolution and magnification objectives. This has allowed us to obtain 100x images of post-expanded samples (Fig. 3c and d, iii and Figs. S14-15).

**Figure 3:**
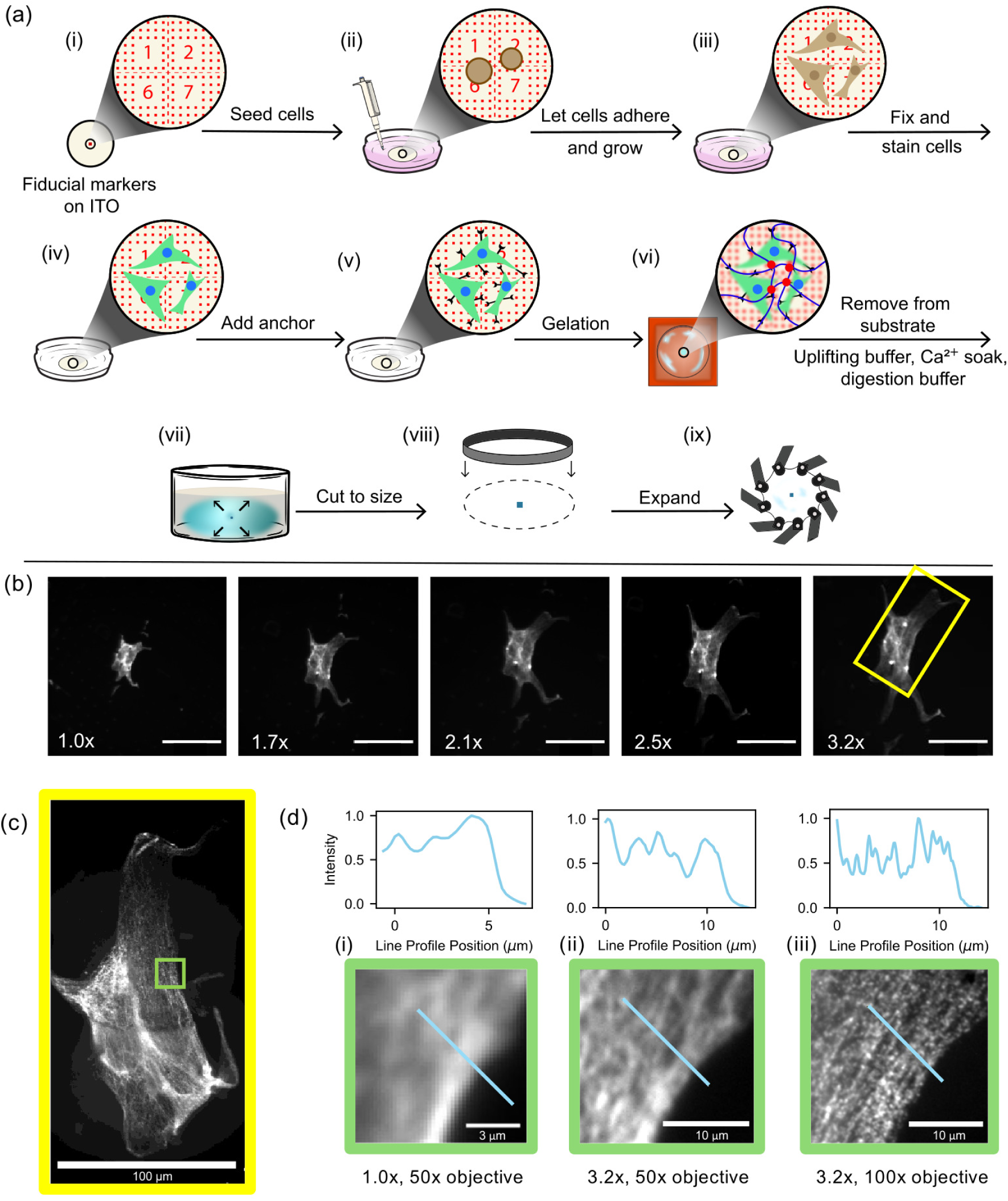
Monitoring fixed cell expansion while applying tensile force to super-resolve microtubular structure. TExM was applied to fixed NIH 3T3 fibroblast cells stained with Alexa 488 for *α*-tubulin. (a) Schematic of workflow for applying TExM to fixed cells. (i) Fiducial markers are printed on ITO coverslips. (ii) Cells are seeded on the ITO coverslips within cell media. (iii) Cells grow and adhere until reaching the desired confluency. (iv) Cells are fixed and stained using a standard immunochemistry approach and (v) are incubated in AcX. (vi) The ITO coverslips with stained, fixed, and AcX treated cells are placed into a silicone-mold and the hydrogel is poured over top. (vii) The ITO- coated glass coverslips are removed from the hydrogel in a series of soaking in uplifting buffer, CaSO_4_, and digestion buffer solutions, leaving the cells embedded within the hydrogel (see Methods). (viii) The hydrogel is cut to size for the iris expansion device with a circular punch cutter. (ix) The hydrogel is expanded on the iris expansion device. (b) Scalebars are 100 *µ*m. Tracking of the same cell during different expansion steps shows expansion using a 50x air objective. Expansion factors correspond to the expected expansion factors. The area outlined in yellow in the last panel is the same area shown in (c) rotated counterclockwise, a 100x oil immersion objective image of the same cell shown after the hydrogel has been dried and sandwiched between glass coverslips. (d) Comparison of resolution of the green outlined area in c) between (i) pre-expansion and (ii) post-expansion of the hydrogel with a 50x objective and (iii) dried hydrogel with 100x objective. The intensities have been normalized to a maximum value of 1. Illustrations in (a) attributed to Allison Breuler, reuse copyright CC BY-NC-ND 4.0.

NIH 3T3 fibroblast cells^28^ were selected for their ordered tubulin structure.^43–45^ Glass coverslips coated in ITO are used as a substrate to allow for cell adhesion and growth in a native manner, as well as clean and easy printing of the fiducial markers based on the refractive index, in addition to clear optical imaging (Fig. 3a, i-iii). Following standard fixation and staining procedures (Fig. 3a, iv), cells are treated with Acryloyl-X (AcX) to cross-link the cells and hydrogel (Fig. 3a, v).^21^ The hydrogel precursor solution is poured onto the treated cells and polymerization is allowed to occur (Fig. 3a, vi). However, the strong -OH interactions between the hydrogel network and ITO surface leads to difficulties in removal of the hydrogel from the substrate by simply peeling. To alleviate the hydrogel/substrate interactions, the sample is treated with an uplifting buffer containing 0.8 M guanidinium hydrochloride, a known chaotropic agent, to disrupt hydrogen bonding between the Alg chains and the ITO-coated substrate. The uplifting buffer induces osmotic swelling, facilitating partial uplifting at the hydrogel periphery. A subsequent CaSO_4_ slurry soak reinforces the hydrogel network, resulting in an increase in hydrogel stiffness. The increased stiffness of the matrix and reduced number of dangling bonds reduces adhesive resistance, minimizing tearing during the final separation between the ITO substrate and hydrogel/cell interface (Fig. 3a, vii). The hydrogel remains structurally sound and easy to work with during these processes. After removal, the hydrogel is treated with a Proteinase K digestion buffer to cleave cellular biomolecules, mechanically homogenizing them, allowing for uniform expansion. Finally, the hydrogel is cut to size, since the uplifting and digestion buffer soak cause swelling within the hydrogel network (Fig. 3a, vii-viii), and mounted to the iris expansion device for microscopic imaging (Fig. 3a, ix).

Cells expand in proportion to the tensile strain applied as shown by imaging the same cell during the expansion process (Fig. 3b). Osmotic ExM is limited to observing only the pre- and final post-expansion result, since the sample is submerged in swelling buffer in a separate petri dish prior to mounting on a glass coverslip to image.^46^ In contrast, the design of the microscope-compatible iris expansion device for TExM allows for continuous observation of the expansion at step sizes as small as 0.003×. Importantly, monitoring cellular expansion allows observation of cellular structures, showing alterations or rearrangement during the expansion process, as anisotropic expansion due to partial digestion is known to occur and can cause false localization of cellular ultrastructures.^12^ The cell in Fig. 3b increases linearly in size from roughly 27 *µm* by 49 *µm* pre-expansion to 49 *µm* by 120 *µm* post-expansion (Table S1). This gives a 1.8× linear expansion along its short axis and 2.4× linear expansion along its long axis. While the entire cell does get larger, the expansion is not purely homogeneous in both directions. The overall expansion of the cell does not correspond to the inputted 3.2× value on the iris expansion device electronics, noting the need for the internal reference fiducial markers to calculate local expansion. Additional images of post-expanded fixed cells are provided as Figure S14, with diameters reaching 200 *µm*. Overall, we confidently observe lateral expansion of fixed cells through tensile force.

TExM of fixed cells achieves super-resolution imaging of microtubules in fibroblasts. As the cell is expanded, the space between the macromolecules increases, and in this case of immune-stained *α*-tubulin, image resolution noticeably increases post-expansion (Fig. 3d). Using a 50× air objective, pre-expansion images exhibit uniform fluorescence throughout the cell, where a cross-sectional line profile fails to resolve features beyond the contrast between the cell’s interior and exterior (Fig. 3d, i). However, following a 3.2× expansion of the same region, a line profile demonstrate the ability to resolve individual tubulin filaments that were previously indistinguishable while imaging with the same 50× air objective (Fig. 3d, ii). Drying the sample for imaging with a high magnification 100x oil immersion objective further increases the resolution, allowing clear identification of individual tubulin filaments (Fig. 3d, iii and c). Quantitative analysis of line sections from microtubules in additional fixed cells demonstrates a super-resolution of approximately 100 nm between filaments (Fig. S15).

### Continuous imaging of live cells while using Tensile Expansion Microscopy (TExM)

TExM allows investigation of living cells during expansion. What is currently lacking from the classical osmotic ExM protocol is a manner to observe live samples due to the requirements of fixation and digestion. By using tensile forces, cells can be seeded upon the hydrogel, then the hydrogel can be expanded whilst the cells remain alive and adhered to the hydrogel network (Fig. 4a). We report our methodology and observations of individual cells and cell clusters under the extreme 1500% areal strain of TExM.

**Figure 4:**
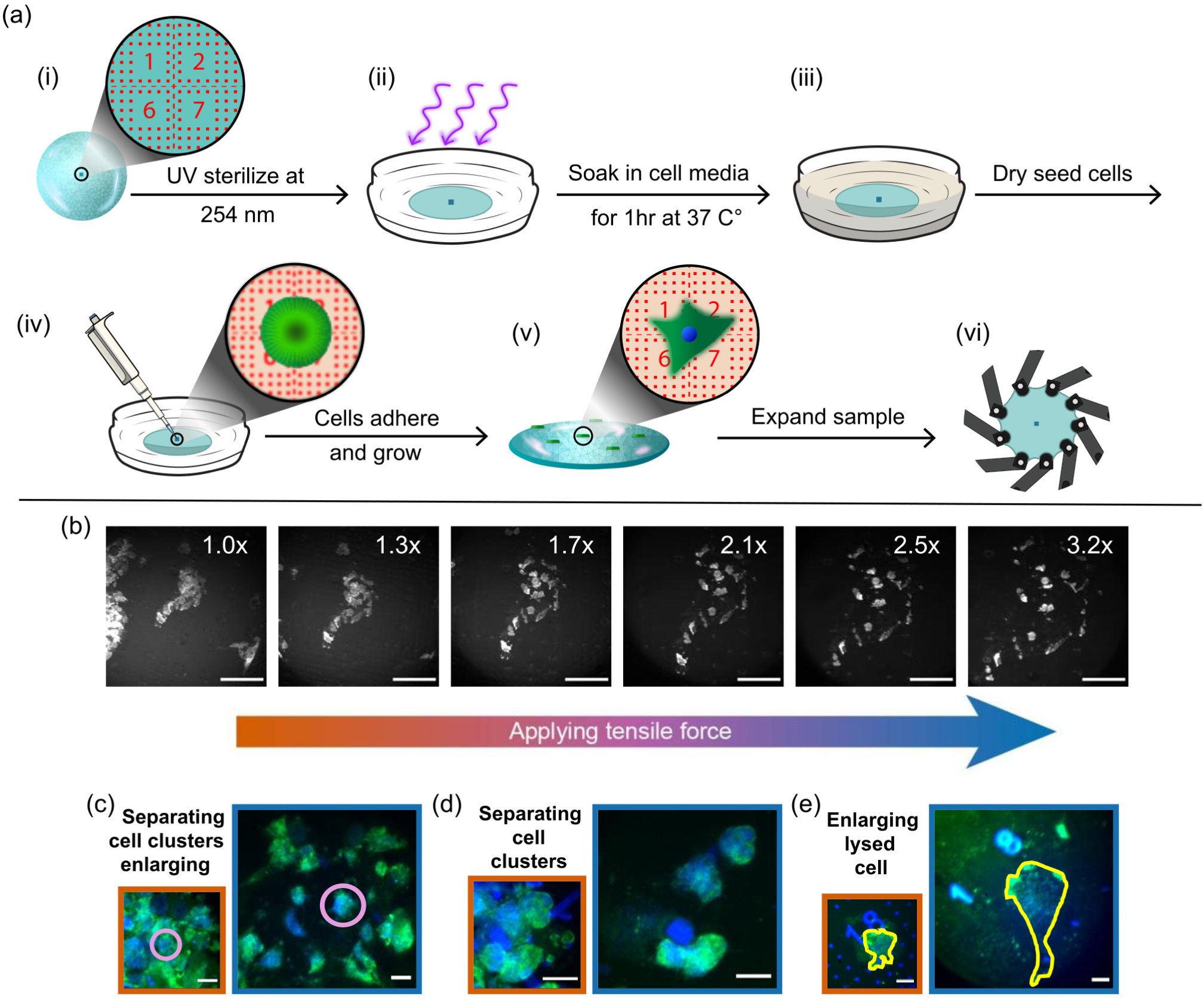
TExM shows increased cell size and distinguishes individual cells through real time imaging of live cell expansion. HeLa cells with microtubule structure (tubulin-mClover3 - green) and soluble cytosolic protein (p62-mRUBY3 - blue) grown on top of a hydrogel. (a) Schematic workflow for applying TExM to live cells. (i) A hydrogel is prepared with fiducial markers embedded. (ii) The hydrogel is sterilized with 254 nm UV light. (iii) The hydrogel is soaked in cell media for 1 hr at 37*^◦^C*. (iv) Cell media is aspirated and a small volume of high density cells in cell media is seeded in the center of the hydrogel and (v) the cells are allowed to grow and adhere. (vi) The hydrogel is expanded on the iris expansion device. (b) Real time observation of a cell cluster separating, but cells and the cluster retain their shape through expansion. Images reported with 488 nm excitation and emission from the tubulin-mClover3. Scalebar is 150 *µm* throughout all images. (c-e) Example behaviors observed of live cells pre- (orange border) and post-tensile expansion (blue border). (c) Clusters are pulled apart and cells increase in size (center cell is circled in purple to assist in visualization of the separation of the cluster). (d) Smaller cell clusters are pulled apart and cells retain their shape. (e) A single cell (outlined in yellow) is lysed and enlarged through out expansion. (c-e) Scalebar is 20 *µm*. Illustrations in (a) attributed to Allison Breuler, reuse copyright CC BY-NC-ND 4.0.

It is important to ensure that live cells adhere to the hydrogels during imaging and expansion. Attempts to functionalize the hydrogel using adhesion proteins (collagen, fibrinogen, poly-lysine) following standard protocols showed NIH 3T3 fibroblast cell adhesion, but led to high cell detachment during hydrogel loading to the iris expansion device.^45,47,48^

This motivated us to switch cell lines to HeLa, due to their robust nature and desire to grow as cell clusters. Hydrogel functionalization was tested for HeLa cells. It was observed that HeLa cells adhere and grow similarly to their native state when the biocompatible DN hydrogel remains non-functionalized. Better expansion behavior occurs on non-functionalized hydrogels compared to those altered with adhesion proteins (Fig. S16). However, care must be taken to ensure cells do not detach from the hydrogel surface when loading into the iris expansion device. For fluorescence visualization, the HeLa cells contain internal tags for microtubule structure (tubulin-mClover3) and soluble cytosolic protein content (p62-mRUBY3). Tubulin-mClover3 and p62-mRUBY3 are represented by green and blue in (Fig. 4c-e), respectively.

TExM can separate living cell clusters and increase individual cell sizes. While the hydrogel expands, several cellular behaviors are observed: cell clusters break apart giving more space between cells (Fig. 4b-d), individual cells moderately expand (Fig. 4c, d) and some cells are shown to lyse (Fig. 4e). Real-time imaging during the expansion (Fig. 4b) showed a group of approximately 25 cells separating. At 1× expansion, the number of cells is difficult to determine, while at first application of tensile force at 1.3×, the cells separate from one another, potentially breaking cell-cell adhesion. With increasing expansion factors, the cells further moved away from one another. In Fig. 4c-e images of the same cells or groups of cells pre- and post-expansion are shown taken with a 50x objective with the hydrogel loaded on the iris expansion device. Both separation and enlargement are observed in Fig. 4c where pre-expansion, it is extremely difficult to determine the cell numbers and boundaries. Post expansion, each individual cell can be clearly identified. To aid the reader in understanding how the dense cell cluster was broken apart, the same cell has been circled in purple, pre- and post-expansion. The size of the purple outlined cell increases by 1.5× linearly, less than the theoretical expansion factor based on the expansion device’s electronics. A cluster of cells is shown in Fig. 4d, where pre-expansion imaging makes it difficult to ascertain the total cell count and the nature of cellular adhesion. Post-expansion imaging shows four distinct cells clearly resolved, two maintaining strong cell-cell adhesion and two that have begun to separate. Finally, Fig. 4e reports a cell, outlined in yellow, that has increased in size by 2.5× linearly, albeit lysing in the process.

The observation of live cells separating and expanding in size provides promise for future application of TExM. Notably, the results in Fig. 4b-e support the ability to monitor expansion and distinguish individual cells. Further evidence can be found in the Fig. S17 where the cells are shown with fiducial markers. Despite cell lysis occurring in some samples, the ability for the iris device to span a range of expansion factors could allow expansion to be stopped at factors where cells are intact and higher resolution is achieved, but prior to larger expansion factors where the cell may lyse and failure occurs (Fig. 4e). Further, we note the current results were obtained under non-ideal conditions, where the experiments were performed at room temperature with low humidity during the expansion process. Imaging in a controlled environment with a microscope-compatible live cell incubator with controlled temperature (37 *^◦^C*), CO_2_, and humidity could lead to higher cell survival. The low humidity further led to dehydration of the hydrogel, limiting the time imaging could be performed. This constraint limits our ability to study expansions at larger magnitudes over extended periods. Consequently, the cellular response to slow, high-strain deformation remains an intriguing area for future investigation. Overall, the ability to separate cells from one another is promising for single cell analysis methods, despite the current live cell expansion factors falling short of super-resolution capabilities to resolve structural components of the cell. TExM can benefit and advance the study of mechanobiology, by allowing much larger perturbations in a controlled manner. By altering the hydrogel substrate through mechanical stimulus, attached cells responses to the perturbation can be observed in real time. However, further controls must be considered while designing experiments and interpreting results due to altering effects of mechanical stimulus on a system.^49^ Future works will test the cellular response to the physiologically-relevant and extreme strains of TExM.

## Conclusion

Tensile Expansion Microscopy (TExM) is an optical imaging method that uses tensile force to physically increase the size and separation of biological features enabling super-resolution imaging of fixed samples, while providing enhanced-resolution monitoring of living cells in two dimensions. TExM relies on four key components: (1) a biocompatible, extensible, and tough hydrogel (Fig. 1a); (2) an expansion device that applies tensile force with coordination of the microscope stage to maintain sample focus during imaging (Fig. 1b); (3) microscale fiducial markers to track sample location and potential local deformations (Fig. 2); and (4) fixed or adherent cell methodologies that do not significantly disrupt hydrogel extensibility properties (Figs. 3a and 4a). Here, we observed precise and repeatable isotropic expansion of up to 4× with local low, submicron variability of expansion at the microscale. Real time tracking of fixed NIH 3T3 fibroblast cells were found to increase in size with the application of tensile force. At the final expanded state, cell diameters reached sizes as large as 200 *µm* and super-resolution imaging was demonstrated through the field-standard measurement of microtubule separation (Figs. S14 and S15). Notably, TExM allowed for the investigation of living cells during expansion, showing diverse behaviors of separating clusters of cells to resolve individual cells and expanding living cells to larger sizes.

The future possibilities for TExM span improving expansion microscopy metrics and applying the method to live cell fluorescence microscopy studies in mechanobiology, other analytical imaging techniques, and even materials science. Two-photon lithography fiducial markers that do not enlarge, fragment, or dilute during expansion, can be fabricated in diverse shapes down to 200 nm and variable stiffness, and can be doped with diverse chemicals. This allows improvement in tracking and quality control in expansion microscopy beyond the current extrinsic references or protein-based markers.^16^ In applications to biophysical studies, the variable 14-16 wt.% hydrogels and iris expansion device provide a versatile tool to vary substrate stiffness and strain at physiological levels for mechanobiology studies, giving access to the extreme tensile strain applied for TExM, previously unavailable. Different cellular behaviors in response to expansion rates were observed and could be further studied. Given the two-dimensional nature of the expansion with the sample thinning axially (Fig. S12), imaging biomacromolecular structures organized at the hydrogel/cell interface, such as focal adhesions, could be pursued in future work. We note that the axial compression could be a limitation to studying tissues and cells where the axial organization is important. The ability of TExM to image expanded cells without fixatives and in dried hydrogels could be applied to increase the spatial resolution of other analytical imaging methods sensitive to immobilizing chemicals and water such as mass spectrometry or Raman microspectroscopy. While expansion mass spectrometry imaging^7^ ^50^ and expansion Raman microscopy^51^ ^52^ methods have been reported based on the osmotic swelling, these studies have mainly focused on tissues or required advanced instrumentation. Finally, the iris expansion device can be used to super-resolve physicochemical phenomenon in the double network hydrogel itself with the inclusion of deformation-sensitive fluorescent reporters.

## Methods

### Hydrogel formulation

#### Preparation of stock solutions

It is highly recommended to use weighing papers instead of plastic or aluminum weighing containers to measure the required weights of all the solid reagents. Weighing paper has limited static effects which improves the accuracy of the formulation. Balance anti-static brushes should further be used to reduce static. Proper balance protocols (e.g., closing doors, balance placement away from vibrations, clean and dry containers) further improve accuracy. The stock solutions homogeneity should be evaluated before use, especially beyond 10–15 days. All samples were prepared with Type I water purified to 18.2 *M* Ω with conductivity lower than 9 *µ*S on an Elga Chorus system unless noted otherwise.

#### Alg stock solution

To prepare 150 mL of a 3% w/v Alg stock solution, 150 g of water was first measured into a glass reagent bottle (Fisherbrand™, Fisher Scientific). Alginic acid sodium salt from brown algae (CAS: 9005-38-3; Sigma Aldrich) was used. Using weighing paper, 4.5 g of the Alg was measured out and was then slowly added to the water while maintaining vigorous magnetic bar stirring at 400–500 rpm. Stirring was crucial to ensure good dispersion of the Alg into the water. Once all the Alg was added, the bottle was closed and sealed with parafilm tape. Stirring was continued vigorously until an increase in the solution’s viscosity was observed. At this point, the stirring velocity was decreased to 50 rpm to avoid shear-induced scission of the Alg chains, which would alter the rheological properties of the precursor solution. The slow mixing was allowed to progress for 1-2 days. Once the solution became completely homogeneous, it was stored at room temperature for no more than 1 week.

#### AAm stock solution

A 50 mL stock solution of 40% w/v acrylamide (AAm) was prepared using acrylamide electrophoresis reagent (CAS: 79-06-1; purity ≥ 99%, Sigma Aldrich). First, 50 mL of water was added to a glass reagent bottle (Fisherbrand™, Fisher Scientific). Next, 20 g of AAm was measured and added to the water. The mixture was stirred vigorously at 300–400 rpm for approximately 1 hour while covered with aluminum sheet to protect it from light, as AAm is susceptible to photodegradation, especially in UV light.^53^ Once the AAm solution became homogeneous, it was immediately used. It is recommended to use freshly prepared AAm stock solutions.

#### MBA stock solution

A 10 mL stock solution of 2% w/v MBA was prepared using N,N’-methylenebisacrylamide (CAS: 110-26-9; purity ≥ 99.5%, Sigma Aldrich). To start, 0.2 g of MBA was added into a 25 mL conical tube (Eppendorf). Then, 10 mL of water was added using a 10 mL pipette, and mixing was performed under vigorous stirring (400 rpm) for about 1 hour using a small stirring bar. Once the MBA solution became homogeneous, it was stored in the fridge at 4 °C. As a precaution, the MBA stock solution was covered with an aluminum sheet to protect it from UV light, as high UV intensity or long UV exposure can lead to the hydrolysis of the MBA amide groups. ^54^

#### APS stock solution

To prepare a 2 mL stock solution of 10% w/v APS, 0.2 g of ammonium persulfate (CAS: 7727-54-0; purity ≥ 98%, Sigma Aldrich) was added to a 25 mL conical tube (Eppendorf). Following this, 2 mL of water was added using a pipette, and the mixture was vortexed for 5 minutes, as APS is very hydrophilic and dissolves rapidly in water. The APS stock solution was freshly prepared before each hydrogel precursor solution preparation and could not be stored.

#### Preparation of Alg-Ca^2+^/PAAm DN hydrogel

Alg-Ca^2+^/ PAAm DN hydrogels were synthesized using two distinct formulations, designated as hydrogel (14 wt.%) and hydrogel (16 wt.%), corresponding to total polymer concentrations of 14 wt.% and 16.4 wt.%, respectively. Across both formulations, the mass ratio of AAm to Alg was maintained at 8.03. The concentration of MBA was fixed at 0.055 wt.% relative to the weight of AAm. Similarly, the concentrations of APS and TEMED (CAS: 110-18-9; purity = 99%, Sigma Aldrich) were each fixed at 0.45 wt.% relative to the weight of AAm. The density of TEMED (0.78 g/cm^3^ at 20°C) was considered in all the calculations of the required volumes.

An example of a 42.68 mL precursor solution preparation of hydrogel is detailed as follows. First, 26.63 g of the Alg stock solution was measured and added to an 80 mL glass reagent bottle with screw cap (Fisherbrand™, Fisher Scientific). The Alg was dispensed from the stock solution using a 30 mL BD Luer-Lok™ plastic sterile syringe without a needle, with the weight precisely controlled on a digital scale. Subsequently, 15.56 mL of the AAm stock solution were added using a volumetric pipette. Following this, 280.1 µL of the APS stock solution and 171.2 µL of the MBA stock solution were added using micropipettes. After the addition of all components, the reagent bottle was sealed with a screw cap, wrapped in an aluminum sheet and gently shaken by hand in a circular motions for 1 minute. The solution was then subjected to magnetic stirring at a low velocity (50-100 rpm) for 45 minutes to ensure homogeneity. The hydrogel (14 wt.%) formulation was prepared following the same procedure as described for hydrogel. However, for hydrogel (14 wt.%), an additional 7.32 mL of water was added to the precursor solution in order to adjust the total polymer concentration to 14 wt.%. A summary of which data were collected with hydrogel (14 wt.%) vs. hydrogel (16.4 wt.%) are presented in Table S2.

Following the 45 minutes stirring period, 36.14 µL of TEMED was added to the solution. The mixture was again gently stirred not shaken by hand for 1 minute before being stirred magnetically at 100 rpm for 15 minutes. Once the precursor solution was homogeneous after TEMED addition, it was poured into the desired mold. A typical molding setup involved a circular silicone mold (50 mm in diameter and 1.5 mm in thickness) placed on a glass plate covered with a crystalline polyethylene terephthalate (PET) thin sheet (Amazon). The mold was then sealed by placing a second glass plate, also covered with a PET sheet, on top of the silicon mold, creating a “sandwich” structure which was critical to minimize the quenching of the free radical polymerization by oxygen in the air.^55^ This assembly was then secured with large office binder clips. The precursor solution was placed in an oven at 35 *^◦^C* for 5 hours to achieve simultaneous free radical polymerization of AAm and covalent crosslinking with MBA, resulting in the formation of the PAAm covalently crosslinked network. The samples were then left to equilibrate overnight at room temperature to ensure complete network stabilization. After this period, the hydrogel was carefully peeled from the mold and immersed in a 20 mM calcium sulfate dihydrate (CaSO_4_ · 2H_2_O, CAS: 10101-41-4, purity = 99% Thermo Fisher Scientific) slurry for 12 minutes yielding the formation of the Alg-Ca2^+^ ionically crosslinked physical network and thus Alg-Ca^2+^/PAAm DN hydrogel.

### Iris control

The iris expansion device can be precisely controlled using both its control box and software-controlled through python. The expansion parameters can be adjusted based on the experiment, with the smallest expansion step size of 0.003x. The expansion speed can be fine tuned for suitability to the sample condition for each experiment where the maximum speed of the iris arm tip is 5 cm/sec. The slowest optimized expansion speed for cells expansion experiment was 0.01 cm/sec. The automated control of the iris expansion device during data acquisition can be achieved by utilizing the pyserial Python library to establish USB serial communication between the Python interface and the iris expansion device.^56^ Pymmcoreplus, an extended python bindings for the C++ micro-manager core, was used to control stage and objective position during image acquisition.^57^

### Substrate cleaning

Substrates (Glass coverslips, Ted Pella, Prod#: 260368 - 30, or Si wafers) were cleaned using a two-step process involving a base solution (TL1) followed by plasma cleaning. First, the TL1 solution was prepared in a fume hood by mixing water, ammonium hydroxide (*NH*_4_*OH* ACS grade, CAS: 1336-21-6; Fisher Scientific) and 30% hydrogen peroxide (*H*_2_*O*_2_ ACS grade, CAS: 7722-84-1; Fisher Scientific) in a 6:1:1 volumetric ratio, respectively. This solution was heated to 80 *^◦^C* using a hotplate equipped with a temperature probe. The substrates were loaded into a holder, dipped in water to remove dust, and then submerged in the 80 *^◦^C* TL1 solution for 90 seconds. Following this, the substrates were rinsed in a second dish of water, dried with ultrapure nitrogen (Airgas), and transferred to a metal sample holder. The metal holder was placed in the plasma cleaner chamber (Harrick Plasma, PDC-32G, 115V), which was then sealed and pumped down using a vacuum pump. To facilitate a high oxygen environment, the chamber was flooded with oxygen for 30–60 seconds and pumped down repeatedly for a total of three cycles. Finally, the chamber pressure was allowed to equilibrate with the oxygen valve open, and the samples were treated with plasma on the “medium” setting for 2 minutes before being vented to atmospheric pressure and stored in a desiccator.

### Two-photon lithography printing of microscale fiducial markers

Substrates (ITO coated glass coverslips, see deposition details later in Methods, or Si wafers) were first cleaned as described in the cleaning substrate methods. Substrates were secured onto appropriate sample holders and loaded into the Nanoscribe Photonic Professional GT2 machine. A suitable two-photon resin (IP-S or IP-Visio) was chosen to print fiducial markers. IP-Visio,^37^ a low auto-fluorescence negative tone photoresist designed for fluorescence microscopy, was doped with 0.1 *mM* Atto-633. IP-S, a negative tone photoresist with high auto-fluorescence in the visible spetra, ^41^ can also be doped with fluorophores to increase fluorescence signal (Fig S9). A drop of resin was applied on the writing objective (25x medium feature set). Using dip-in mode, the patterns were printed on the wafer through a two-photon process (Fig. 2 a,i). Printing parameters used for each resin are noted in Table 1. The features were developed in propylene glycol methyl ether acetate (PGMEA; CAS: 108-65-6, Sigma Aldrich) for 20 minutes and cleaned in isopropyl alcohol (IPA; CAS: 67-63-0, Sigma Aldrich) for 5 minutes. The patterned substrates were then dried using a nitrogen gun and stored in the dark until ready to use.

**Table 1:** Two photon printing parameters.

| | Power Scaling | Laser Power (%) | Scan Speed ($\mu m/s$) | Galvo Acceleration ($V/ms^2$) |
| --- | --- | --- | --- | --- |
| IP-Visio | 1 | 80 | 2000 | 0.4 |
| IP-S | 1 | 100 | 10000 | 1 |

### Transfer of microscale fiducial markers from Si wafer into hydrogels

A mold was prepared as shown in Fig. 2a, iii with a silicone spacer (50 mm in diameter and 1.5 mm in thickness) on a Si wafer with fiducial markers. Hydrogel precursor solution was poured into the mold and sealed. AAm was polymerized and covalently crosslinked resulting in PAAm chemical network as described in the hydrogel formulation section. The polymerized hydrogel over the Si wafer was soaked in 20 mM CaSO_4_ · 2H_2_O slurry for 6 minutes. This step is crucial to increase the stiffness of the hydrogel, resulting in a hydrogel less prone to tearing while peeling from the Si wafer. The hydrogel was then gently peeled by hand from the Si wafer and re-soaked in the CaSO_4_ · 2H_2_O slurry for an additional 6 minutes to complete the formation of Alg-Ca^2+^ physical network.

### Fluorescence microscopy

Imaging was performed on an inverted fluorescence wide field microscope that has been previously described^58^ with additional lasers and brightfield module. The base of the microscope consisted of an Olympus IX-83 microscope body and brightfield light pillar with halogen lamp (IX3-ILL, U-LH100-3-7). The home-built super-resolution fluorescence microscopy setup used free-standing optics for excitation by either a 100 mW 488 nm (Toptica iBeam Smart), 100 mW 561 nm (CNI), or 100 mW 633 nm (Toptica iBeam Smart) laser. The maximum power output of the laser beams were attenuated using neutral density filters (ThorLabs) or are controlled using Toptica software. The beams were expanded to a diameter of 7.5 mm and then eventually to 22.5 mm passing through a lens (focal length = 400 mm) that focused the beam to the center of the back aperture of the objective lens. The dichroic mirrors (either Chroma ZET488rdc, ZET561rdc, or ZET633rdc for 488, 561, or 633 nm excitation, respectively) then directed the laser beam through either a 20x/0.5 NA air (Olympus UPLFLN), 50x/0.45 NA air (Olympus SLMPlan), or 100x/1.49 NA (Olympus UApo N) oil objective lens in epifluorescence mode. The laser power density at the sample plane ranged from 1.0 – 5.0 mW/cm^2^ depending on the sample. The location of the sample relative to the objective in xyz was controlled via xy piezostage (Olympus IX3-SSU) and objective positioner (Olympus IX3-D6RES). The same objective lens was then used to collect the fluorescence signal from the sample. The emitted light passed through additional emission filters (Chroma ZET488NF, ZET561NF, ZET635NF for 488, 561, or 633 nm excitation, respectively) to remove any stray reflected excitation light. A Photometrics Prime 95B 22 mm scientific CMOS camera was used to capture the 16-bit images at 150 ms exposure times for all cells-related images (Fig. 3 and 4) and 20 ms for fiducial markers images (Fig. 2). For each cell-related image acquisition, where 20 frames were taken and summed together to obtain final images. Camera and microscope stages were controlled using the Micromanager 2.0 open source microscopy software. For the autofocus and multiple field of view acquisition, a developed Python script was used to control all the hardware. The Python pymmcore library controlled microscope stage and camera while the pyserial library controlled the iris expansion device.

### Autofocus Python control of microscope and data collection during expansion

A schematic of automated data acquisition is shown in Fig. 2b. A Python library, Pycro-manager, allowed for binding between Micro-Manager’s MMCore written in Java and Python for the microscope stage movement control and camera control for image acquisition.^59^ The algorithm controled the z-position of the microscope objective relative to the hydrogel to maintain an in-focus image, the collection and storage of images on the microscope camera. Our software autofocus algorithm uses the Tenengrad algorithm, a Sobel gradient-based method known for fast computational speed. The Sobel operator detects intensity variations along both horizontal and vertical directions within the data of the microscale fluorescent fiducial markers or cells on the microscope camera, based on the assumption that sharper images exhibit stronger edges, resulting in higher gradient values. Image sharpness was quantified as scores by computing the mean gradient magnitude across all pixels^60^.^61^ Focus scores for each images was acquired from z-scan of 3 *µm* step sizes (resolution), the highest focus score indicated the in-focus image. The Tenangrad formula is as follows:

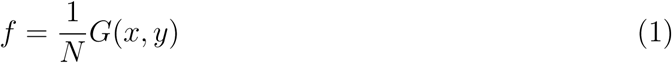

where N is the total number of pixels and the expression of *G*(*x, y*) is:

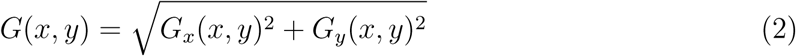

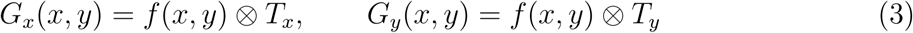

The operator ⊗ represents the convolution operation. The gradient operators *T_x_* and *T_y_* are defined such that *T_y_* is the transpose of the matrix 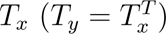, expressed as:

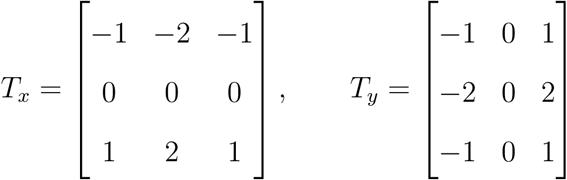

### Multiple field of view acquisition and image stitching

A custom Python function was developed to acquire multiple fields of view (FOVs) centered around a selected object. To generate a N x N grid pattern around the feature of interest, the physical offset between adjacent FOVs was calculated as:

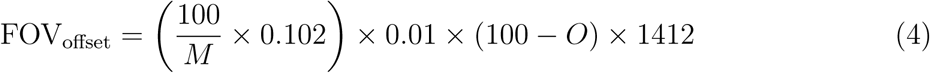

where M is the objective magnification, the pixel size corresponding with our 100x magnification objective is 0.102 *µ*m, *O* is the desired overlap percentage, and 1412 corresponds to the camera sensor width in pixels. The offset determines the lateral displacement between consecutive image positions in micrometers. At each grid position, the xy stage was moved accordingly, and the autofocus algorithm described previously (see autofocus python control methods) was executed utilizing a coarse focus search with a bidirectional scan of 20 *µ*m at a step size of 1 *µ*m followed by a fine focus search starting at the z-position corresponding to the peak focus score for the coarse focus search. The fine search is a bidirectional scan of 5 *µ*m at a 0.25 *µ*m step size. The in focus image is obtained at the z-position corresponding to the highest focus score calculated during the fine focus search. All acquisitions are started at a near in focus position set manually. Due to hardware limitations, like mechanical stage repeatability and camera alignment angle With respect to the microscope stage, the grid of images cannot be stitched naively to get a well aligned larger field of view. Microscopy Image Stitching Tool (MIST)^62^ is used to account for the hardware limitations and use software optimized for microscopy imaging to obtain stitched images used for further analysis. MIST software was acquired as an ImageJ plugin from the National Institute of Standards and Technology.

### Image registration of microscale fluorescent fiducial markers

The pre- and post-expansion stitched images were individually binarized using global thresholds. The threshold was chosen such that a maximum number of fluorescent features can be identified without distortion to their shape. A MATLAB function, regionprops,^63^ was used to find unique objects in the binarized images and calculate their centroids. The objects were filtered on the basis of their area, keeping only those objects that correspond to the fluorescent fiducial markers. Objects representing the fiducial markers on the pre- and post-expansion images were correlated manually with care taken to ignore objects whose centroids do not correspond to any fiducial markers. MATLAB’s inbuilt image registration estimator (estgeotform2d)^64^ was used to calculate an optimal similarity transformation between the pre-expansion and post-expansion image. The transform was estimated 50 times. Each transform corresponds to a scaling, translation and rotation estimate. The expansion factor was obtained by averaging the corresponding 50 expansion factors. The mismatch between the objects post transformation was taken to be real deformation at that location. A vector deformation field was calculated using a triangulation based linear interpolation method, similar to the methods used in the image registration of macroscale fiducial markers.^65^ A Root Mean Square error was also calculated based on the vector deformation field within the convex hull of the centroids of identified fiducial markers

### Image registration of macroscale fiducial markers

The macroscale fiducial markers were identified using MATLAB’s inbuilt blob detection algorithm^63^ and were tracked through the video using a nearest neighbor algorithm. MATLAB’s inbuilt image registration estimator (estgeotform2d)^64^ was used to calculate an optimal similarity transformation between the pre-expansion (Fig. 1c) and post-expansion (Fig. 1d) image. A deformation field (red arrows Fig. 1e) was estimated by evaluating the difference between the registered post-expansion image and the pre-expansion image by using a triangulation based linear interpolation method. A scaling factor was obtained from the rigid registration, which was the expected expansion factor. See SI method and Figs. S3-5 for more details.

### Fixed cells in hydrogels

#### ITO coating of glass coverslip

Coverslip glass (Ted Pella, Prod#: 260368 - 30) was cleaned as described in the substrate cleaning methods. 100 nm of 95/5 ITO was DC magnetron sputtered using the Angstrom EvoVac Multiprocess Physical Vapor Deposition (PVD) System. After ITO coating, fluorescent fiducial markers are printed on the coverslip glass as described in the two-photon lithography methods section.

#### Cell plating for fixed cells

ITO coated glass coverslips, with fluorescent fiducial markers, were sterilized with 254 nm UV for 1 hour, then flipped to the alternate side and exposed for another hour. Each sterilized coverslip (Ted Pella, Prod#: 260368 - 30) with printed fiducial markers was covered with 1.5 mL of complete growth medium [Dulbecco’s Modified Eagle Medium (DMEM) (SH30022.01; Cytiva) supplemented with 10% fetal bovine serum (FBS) (SH30071.03; Cytiva)] in a six well plate (657 160; CellStar). NIH 3T3 rd12 cells (citation) were suspended in complete growth medium. Then 0.5 mL of cell suspension was added dropwise directly into each well. 3T3 cells were placed in a humidified incubator at 37 *^◦^C* with 5% CO_2_ and maintained until cells reached near 80% confluency.

#### Fixation and immunology

Cells were washed three times with 1 mL Dulbecco’s phosphate-buffered saline (DPBS) (14190-144; Gibco) and fixed with warmed (37 *^◦^C*) 4% paraformaldehyde (15710; Electron Microscopy Sciences) in DPBS for 10 min at room temperature. Following fixation, paraformaldehyde was aspirated and cells were rinsed three times with DPBS (1 mL per well). Cells were permeabilized with 0.1% Triton X-100 (TX-100, A16046.AE; ThermoScientific) in DPBS for 15 min at room temperature, followed by three rinses with DPBS. Cells were then incubated overnight at 4*^◦^C* with anti-*α*-tubulin antibody Alexa Fluor 488 (53-4502-82; Invitrogen) (5 *µ*g/mL with 0.1% BSA (126593-10GM; EMD Millipore) in DPBS). Cells were rinsed three times with 1 mL DPBS and incubated with secondary antibody (A21121; invitrogen) (1 *µ*g/mL with 0.1% BSA in DPBS) for 2 hr at room temperature. After staining, cells were washed three times with DPBS.

#### Acryloyl-X (AcX) anchoring treatment

To enable subsequent hydrogel embedding, cells were incubated with 0.1 mg/mL Acryloyl-X (AcX) (A20770; Invitrogen) in DPBS for 3 h at room temperature. Following incubation, cells were washed twice with DPBS for 15 min each.

#### Polymerization of hydrogel over fixed cells

Fixed 3T3 cells on ITO-coated glass coverslips with printed fiducial markers were positioned within circular silicone molds (28 mm diameter, 1.5 mm thickness). Once the coverslips and molds were prepared, TEMED was added as the final reagent to the hydrogel precursor solution in a separate reagent bottle. The mixture was homogenized via gentle manual agitation for 1 min, followed by 15 min of magnetic stirring, then poured onto the pre-prepared molds. The molds were sealed with PET sheets and glass plates, secured with binder clips, and incubated at 35*^◦^C* for 5 hours to ensure complete AAm polymerization and covalent crosslinking. Finally, the hydrogels were allowed to equilibrate overnight at room temperature.

#### Substrate removal

Following the formation of the PAAm covalently bonded network, the coverslips with the adhered hydrogels were removed from the molds. The hydrogels were then placed in a 6-well plate containing a detachment buffer composed of 0.8 M guanidine hydrochloride (GuHCl; CAS: 50-01-1; Sigma-Aldrich), 50 mM tris(hydroxymethyl)aminomethane (Tris base titrated to pH = 8.0 with HCl; CAS: 77-86-1; purity ≥ 99.9%, Promega), and 0.5% v/v TX-100. This step is essential to initiate the uplifting of the hydrogel from the ITO-coated substrate. The plates were then incubated at 37*^◦^C* under orbital shaking of 120 rpm for 3 hours to initiate the uplifting on the sides. Subsequently, the assemblies were immersed in a 20 mM CaSO_4_ · 2H_2_O slurry for 6 min to initiate Ca^2+^-mediated ionic crosslinking of Alg. The remaining adhered region was then detached from the ITO-coated glass coverslip using a controlled, solvent-assisted peeling technique. Given the mechanical compliance of the hydrogel relative to the substrate adhesion, the detachment process required careful manual handling and a quasi-static peeling rate to preserve sample integrity. To prevent cohesive failure (tearing), the peeling front was advanced slowly. Upon observing resistance or localized pinning, the propagation was stopped, and water droplets were introduced directly at the crack tip. Consistent with the capillary peeling mechanism, the solvent was allowed to wick into the interface, reducing the effective fracture energy.^66^ Crucially, this step necessitated a patient, iterative approach, allowing sufficient equilibration time for the solvent meniscus to weaken the adhesion before peeling resumed. This slow, stepwise propagation of the fracture front ensured that the applied stress remained consistently below the hydrogel’s bulk cohesive strength, resulting in a damage-free, clean adhesive failure. Immediately after detachment, the hydrogels were re-immersed in the 20 mM CaSO_4_ · 2H_2_O slurry for an additional 6 min to ensure complete bilateral crosslinking of the Alg network from both surfaces. Finally, the detached hydrogels and the original ITO-coated coverslips were inspected under a microscope to evaluate the transfer efficiency of the 3T3 cells and fiducial markers, and to identify regions of interest (ROIs) for subsequent targeting during tensile expansion microscopy.

#### Digestion and homogenization

Following the secondary soaking step in CaSO_4_ · 2H_2_O slurry, the hydrogels were thoroughly rinsed with water to remove excess ions. Then, to achieve isotropic expansion, the samples were transferred to a 6-well plate containing a digestion buffer composed of 50 mM Tris (pH 8.0), 0.5% v/v TX-100, and 0.3 mg/mL Proteinase K (CAS: 39450-01-6; Thermo Fisher Scientific). The TX-100 serves to permeabilize the cellular membranes, ensuring full enzymatic access to the intracellular space.^67^ The samples were incubated at room temperature for 1 hour. In accordance with conventional ExM protocols, this proteolytic digestion step is necessary to cleave peptide bonds within the fixed 3T3 cellular architecture, thereby homogenizing the mechanical properties of the biological specimen. This process is critical to resolve the mechanical mismatch between the stiff cellular network and the swollen hydrogel, ensuring that the subsequent expansion is uniform and distortion-free. Following the 1 hour enzymatic digestion, the hydrogels were removed from the digestion buffer and rinsed thoroughly with water to arrest proteolytic activity and remove residual debris. To ensure precise compatibility with the optical setup, the hydrogels were then trimmed to a standardized 32 mm diameter using a circular punch cutter, preparing them for subsequent mounting onto the imaging iris.

### Live cells on hydrogels

#### Sterilization of hydrogels

Hydrogels were sterilized by ultraviolet (UV) irradiation^68^ using a 254 nm UV source (Amazon). Samples were exposed to UV for 20 min, rested for 10 min, and then re-exposed for an additional 20 min. This sterilization protocol was applied consistently to all hydrogels.

#### Cell seeding

Hydrogels were incubated in 3 mL of complete growth medium DMEM supplemented with 10% FBS for 30 min at 37*^◦^C* and 5% CO_2_ to equilibrate prior to cell seeding. Growth medium was aspirated and hydrogels were allowed to dry for 30 minutes under sterile conditions before use. HeLa cells were seeded onto the air dried hydrogels at 1 million cells per mL in 250 microlitters of culture medium. Hydrogels were then incubated at 37*^◦^C* with 5% CO_2_, and left overnight for cell adhesion.

#### Loading onto iris expansion device

The hydrogel with adhered, live cells is carefully loaded into the iris device with the help of gel loading paraphernalia (Fig. S2). Care is taken to ensure cells are not removed from the hydrogel surface.

## Acknowledgement

This research was financially supported by an Allen Distinguished Investigator Award supported by Allen Family Philanthropies, a Scialog program sponsored jointly by Research Corporation for Science Advancement and the Gordon and Betty Moore Foundation and includes grant number 28410, and NIH NIGMS R35GM142466. We also thank Professor Peng Zhang for guidance in osmotic expansion microscopy, Professor Yi Zhang and Dr. Dechen Fu for providing fluorescently labeled HeLa cells, and Professor Adrian Ranga for insight on the iris design. The NIH 3T3 rd12 cells were a gift from Dr. Krzysztof Palczewski of UC Irvine. We also appreciate Cole Reinholt, Zhanda Chen, and Andrew Maytin for their early contributions to this project, Dr. Zechariah Pfaffenberger for consultation on the microscope, Dr. Andrew Liniger of the Materials for Opto/Electronics Research and Education Center for support with two-photon lithography, members of the Andresen Eguiluz, Mathur, and Sanchez labs for useful discussions, and the entire Kisley lab for discussion.

## References

(1) Chen, F.; Tillberg, P. W.; Boyden, E. S. Expansion microscopy. Science 2015, 347, 543–548.

(2) Yuan, P.; Zhang, M.; Tong, L.; Morse, T. M.; McDougal, R. A.; Ding, H.; Chan, D.; Cai, Y.; Grutzendler, J. PLD3 affects axonal spheroids and network defects in Alzheimer’s disease. Nature 2022, 612, 328–337.

(3) Gao, R.; Asano, S. M.; Upadhyayula, S.; Pisarev, I.; Milkie, D. E.; Liu, T.-L.; Singh, V.; Graves, A.; Huynh, G. H.; Zhao, Y.; others Cortical column and whole-brain imaging with molecular contrast and nanoscale resolution. Science 2019, 363, eaau8302.

(4) Chen, F.; Wassie, A. T.; Cote, A. J.; Sinha, A.; Alon, S.; Asano, S.; Daugharthy, E. R.; Chang, J.-B.; Marblestone, A.; Church, G. M.; others Nanoscale imaging of RNA with expansion microscopy. Nature Methods 2016, 13, 679–684.

(5) Wang, Y.; Eddison, M.; Fleishman, G.; Weigert, M.; Xu, S.; Wang, T.; Rokicki, K.; Goina, C.; Henry, F. E.; Lemire, A. L.; others EASI-FISH for thick tissue defines lateral hypothalamus spatio-molecular organization. Cell 2021, 184, 6361–6377.

(6) Alon, S.; Goodwin, D. R.; Sinha, A.; Wassie, A. T.; Chen, F.; Daugharthy, E. R.; Bando, Y.; Kajita, A.; Xue, A. G.; Marrett, K.; others Expansion sequencing: Spatially precise in situ transcriptomics in intact biological systems. Science 2021, 371, eaax2656.

(7) Zhang, H.; Ding, L.; Hu, A.; Shi, X.; Huang, P.; Lu, H.; Tillberg, P. W.; Wang, M. C.; Li, L. TEMI: tissue-expansion mass-spectrometry imaging. Nature Methods 2025, 22, 1051–1058.

(8) Asano, S. M.; Gao, R.; Wassie, A. T.; Tillberg, P. W.; Chen, F.; Boyden, E. S. Expansion microscopy: protocols for imaging proteins and RNA in cells and tissues. Current Protocols in Cell Biology 2018, 80, e56.

(9) Wassie, A. T.; Zhao, Y.; Boyden, E. S. Expansion microscopy: principles and uses in biological research. Nature Methods 2019, 16, 33–41.

(10) Chang, J.-B.; Chen, F.; Yoon, Y.-G.; Jung, E. E.; Babcock, H.; Kang, J. S.; Asano, S.; Suk, H.-J.; Pak, N.; Tillberg, P. W.; others Iterative expansion microscopy. Nature Methods 2017, 14, 593–599.

(11) Nakamoto, M. L.; Forró, C.; Zhang, W.; Tsai, C.-T.; Cui, B. Expansion microscopy for imaging the cell–material interface. ACS Nano 2022, 16, 7559–7571.

(12) Guichard, P.; Hamel, V. Expansion Microscopy for Cell Biology; Academic Press, 2021; Vol. 161.

(13) Chozinski, T. J.; Halpern, A. R.; Okawa, H.; Kim, H.-J.; Tremel, G. J.; Wong, R. O. L.; Vaughan, J. C. Expansion microscopy with conventional antibodies and fluorescent proteins. Nature Methods 2016, 13, 485–488.

(14) Damstra, H. G.; Mohar, B.; Eddison, M.; Akhmanova, A.; Kapitein, L. C.; Tillberg, P. W. Visualizing cellular and tissue ultrastructure using Ten-fold Robust Expansion Microscopy (TREx). eLife 2022, 11, e73775.

(15) Gao, R.; Yu, C.-C.; Gao, L.; Piatkevich, K. D.; Neve, R. L.; Munro, J. B.; Upadhyayula, S.; Boyden, E. S. A highly homogeneous polymer composed of tetrahedron-like monomers for high-isotropy expansion microscopy. Nature Nanotechnology 2021, 16, 698–707.

(16) Damstra, H. G.; Passmore, J. B.; Serweta, A. K.; Koutlas, I.; Burute, M.; Meye, F. J.; Akhmanova, A.; Kapitein, L. C. GelMap: intrinsic calibration and deformation mapping for expansion microscopy. Nature Methods 2023, 20, 1573–1580.

(17) Aleksejenko, N.; Heller, J. P. Super-resolution imaging to reveal the nanostructure of tripartite synapses. Neuronal Signaling 2021, 5, NS20210003.

(18) Abberior Expansion microscopy: bigger samples for better resolution. 2024; https://abberior.rocks/knowledge-base/expansion-microscopy-bigger-samples-for-better-resolution/, Accessed: February 19, 2026.

(19) Wang, S.; Shin, T. W.; Yoder, H. B.; McMillan, R. B.; Su, H.; Liu, Y.; Zhang, C.; Leung, K. S.; Yin, P.; Kiessling, L. L.; others Single-shot 20-fold expansion microscopy. Nature Methods 2024, 21, 2128–2134.

(20) Guo, J.; Yang, H.; Lu, C.; Cui, D.; Zhao, M.; Li, C.; Chen, W.; Yang, Q.; Li, Z.; Chen, M.; others BOOST: a robust ten-fold expansion method on hour-scale. Nature Communications 2025, 16, 2107.

(21) Tillberg, P. W.; Chen, F.; Piatkevich, K. D.; Zhao, Y.; Yu, C.-C.; English, B. P.; Gao, L.; Martorell, A.; Suk, H.-J.; Yoshida, F.; others Protein-retention expansion microscopy of cells and tissues labeled using standard fluorescent proteins and antibodies. Nature Biotechnology 2016, 34, 987–992.

(22) Irgen-Gioro, S.; Yoshida, S.; Walling, V.; Chong, S. Fixation can change the appearance of phase separation in living cells. elife 2022, 11, e79903.

(23) Betzig, E.; Patterson, G. H.; Sougrat, R.; Lindwasser, O. W.; Olenych, S.; Bonifacino, J. S.; Davidson, M. W.; Lippincott-Schwartz, J.; Hess, H. F. Imaging intracellular fluorescent proteins at nanometer resolution. Science 2006, 313, 1642–1645.

(24) Balzarotti, F.; Eilers, Y.; Gwosch, K. C.; Gynnå, A. H.; Westphal, V.; Stefani, F. D.; Elf, J.; Hell, S. W. Nanometer resolution imaging and tracking of fluorescent molecules with minimal photon fluxes. Science 2017, 355, 606–612.

(25) Chatterjee, S.; Kramer, S. N.; Wellnitz, B.; Kim, A.; Kisley, L. Spatially resolving size effects on diffusivity in nanoporous extracellular matrix-like materials with fluorescence correlation spectroscopy super-resolution optical fluctuation imaging. The Journal of Physical Chemistry B 2023, 127, 4430–4440.

(26) Deschout, H.; Lukes, T.; Sharipov, A.; Szlag, D.; Feletti, L.; Vandenberg, W.; Dedecker, P.; Hofkens, J.; Leutenegger, M.; Lasser, T.; others Complementarity of PALM and SOFI for super-resolution live-cell imaging of focal adhesions. Nature Communications 2016, 7, 13693.

(27) Sun, J.-Y.; Zhao, X.; Illeperuma, W. R.; Chaudhuri, O.; Oh, K. H.; Mooney, D. J.; Vlassak, J. J.; Suo, Z. Highly stretchable and tough hydrogels. Nature 2012, 489, 133–136.

(28) Ho-lubowicz, R. et al. Safer and efficient base editing and prime editing via ribonucleoproteins delivered through optimized lipid-nanoparticle formulations. Nature Biomedical Engineering 2024, 9, 57–78.

(29) Rowley, J. A.; Madlambayan, G.; Mooney, D. J. Alginate hydrogels as synthetic extra-cellular matrix materials. Biomaterials 1999, 20, 45–53.

(30) Tse, J. R.; Engler, A. J. Preparation of hydrogel substrates with tunable mechanical properties. Current Protocols in Cell Biology 2010, 47, 10–16.

(31) Truckenbrodt, S.; Maidorn, M.; Crzan, D.; Wildhagen, H.; Kabatas, S.; Rizzoli, S. O. X10 expansion microscopy enables 25-nm resolution on conventional microscopes. The EMBO Reports 2018, 19, EMBR201845836.

(32) Mandal, J.; Zhang, K.; Spencer, N. D.; others Oxygen inhibition of free-radical polymerization is the dominant mechanism behind the “mold effect” on hydrogels. Soft Matter 2021, 17, 6394–6403.

(33) Abdel Fattah, A. R.; Daza, B.; Rustandi, G.; Berrocal-Rubio, M. Á.; Gorissen, B.; Poovathingal, S.; Davie, K.; Barrasa-Fano, J.; Cóndor, M.; Cao, X.; others Actuation enhances patterning in human neural tube organoids. Nature Communications 2021, 12, 3192.

(34) Schürmann, S.; Wagner, S.; Herlitze, S.; Fischer, C.; Gumbrecht, S.; Wirth-Hücking, A.; Prölß, G.; Lautscham, L.; Fabry, B.; Goldmann, W.; others The IsoStretcher: an isotropic cell stretch device to study mechanical biosensor pathways in living cells. Biosensors and Bioelectronics 2016, 81, 363–372.

(35) Bighi, B.; Ragazzini, G.; Gallerani, A.; Mescola, A.; Scagliarini, C.; Zannini, C.; Marcuzzi, M.; Olivi, E.; Cavallini, C.; Tassinari, R.; others Cell stretching devices integrated with live cell imaging: a powerful approach to study how cells react to mechanical cues. Progress in Biomedical Engineering 2024, 7, 012005.

(36) Nanoscribe GmbH Definite Focus Interface Finder. https://support.nanoscribe.com/hc/en-gb/articles/360003535714-Definite-Focus-Interface-Finder, Accessed: 2026-02-18.

(37) Nanoscribe GmbH IP-Visio: Technical Data and Application Guide. https://support.nanoscribe.com/hc/en-gb/articles/360011709499-IP-Visio, 2024; Accessed: 2026-02-18.

(38) Liu, H. J.; Kang, R. K.; Gao, S.; Zhou, P.; Tong, Y.; Guo, D. M. Development of a measuring equipment for silicon wafer warp. Advanced Materials Research 2013, 797, 561–565.

(39) Wang, M.; Huang, Y.; Liu, S.; Li, L.; Li, P.; Li, J.; Yan, K. A novel high-precision measurement method for warpage and bow of doubile-side polished wafers based on wavelength-tunable laser interferometry. Measurement 2026, 258, 119124.

(40) Khatua, S.; Godoy, J.; Tour, J. M.; Link, S. Influence of the Substrate on the Mobility of Individual Nanocars. The Journal of Physical Chemistry Letters 2010, 1, 3288–3291.

(41) Nanoscribe GmbH IP-S: Technical Data and Application Guide. https://support.nanoscribe.com/hc/en-gb/articles/360001750353-IP-S, 2024; Accessed: 2026-02-18.

(42) Klimas, A.; Gallagher, B. R.; Wijesekara, P.; Fekir, S.; DiBernardo, E. F.; Cheng, Z.; Stolz, D. B.; Cambi, F.; Watkins, S. C.; Brody, S. L.; others Magnify is a universal molecular anchoring strategy for expansion microscopy. Nature Biotechnology 2023, 41, 858–869.

(43) Bhatia, S. N.; Yarmush, M. L.; Toner, M. Controlling cell interactions by micropatterning in co-cultures: Hepatocytes and 3T3 fibroblasts. Journal of Biomedical Materials Research 1997, 34, 189–199.

(44) Hennig, K.; Wang, I.; Moreau, P.; Valon, L.; DeBeco, S.; Coppey, M.; Miroshnikova, Y.; Albiges-Rizo, C.; Favard, C.; Voituriez, R.; others Stick-slip dynamics of cell adhesion triggers spontaneous symmetry breaking and directional migration of mesenchymal cells on one-dimensional lines. Science Advances 2020, 6, eaau5670.

(45) Wawrzyk, D.; Goncerz, P.; Rajfur, Z. Methodological study of fibroblasts MEF 3T3 heterogeneity cultured on hydrogel substrates with different surface functionalization. bioRxiv, 2025; Preprint available at 10.1101/2025.06.12.659070.

(46) Zhang, C.; Kang, J. S.; Asano, S. M.; Gao, R.; Boyden, E. S. Expansion microscopy for beginners: visualizing microtubules in expanded cultured HeLa cells. Current Protocols in Neuroscience 2020, 92, e96.

(47) Andresen Eguiluz, R. C.; Kaylan, K. B.; Underhill, G. H.; Leckband, D. E. Substrate stiffness and VE-cadherin mechano-transduction coordinate to regulate endothelial monolayer integrity. Biomaterials 2017, 140, 45–57.

(48) Prahl, L. S.; Porter, C. M.; Liu, J.; Viola, J. M.; Hughes, A. J. Independent control over cell patterning and adhesion on hydrogel substrates for tissue interface mechanobiology. iScience 2023, 26, 106657.

(49) Dan, A.; Huang, R. B.; Leckband, D. E. Dynamic imaging reveals coordinate effects of cyclic stretch and substrate stiffness on endothelial integrity. Annals of Biomedical Engineering 2016, 44, 3655–3667.

(50) Chan, Y. H.; Pathmasiri, K. C.; Pierre-Jacques, D.; Hibbard, M. C.; Tao, N.; Fischer, J. L.; Yang, E.; Cologna, S. M.; Gao, R. Gel-assisted mass spectrometry imaging enables sub-micrometer spatial lipidomics. Nature Communications 2024, 15, 5036.

(51) Artur, C. G.; Womack, T.; Zhao, F.; Eriksen, J. L.; Mayerich, D.; Shih, W.-C. Plasmonic nanoparticle-based expansion microscopy with surface-enhanced Raman and dark-field spectroscopic imaging. Biomedical Optics Express 2018, 9, 603.

(52) Shi, L.; Klimas, A.; Gallagher, B.; Cheng, Z.; Fu, F.; Wijesekara, P.; Miao, Y.; Ren, X.; Zhao, Y.; Min, W. Super-resolution vibrational imaging using expansion stimulated Raman scattering microscopy. Advanced Science 2022, 9, 2200315.

(53) Marcus J. Caulfield, G. G. Q.; Solomon, D. H. Some Aspects of the Properties and Degradation of Polyacrylamides. Chemical Reviews 2002, 102, 3067–3084.

(54) Xinming Li, Y. C. Ultraviolet-Induced Decomposition of Acrylic Acid-Based Superabsorbent Hydrogels Crosslinked with N,N-Methylenebisacrylamide. Journal of Applied Polymer Science 2008, 108, 3435–3441.

(55) Norman, R. X.; Chen, Y.-C.; Recchia, E. E.; Loi, J.; Rosemarie, Q.; Lesko, S. L.; Patel, S.; Sherer, N.; Takaku, M.; Burkard, M. E.; others One step 4× and 12× 3D-ExM enables robust super-resolution microscopy of nanoscale cellular structures. Journal of Cell Biology 2024, 224, e202407116.

(56) Liechti, C. pySerial Documentation. version 3.5, 2020; Accessed: 2026-02-16.

(57) Talley, N.; Lambert, T.; pymmcore-plus contributors pymmcore-plus: A pythonic wrapper for micro-manager/mmcore. 2024; https://github.com/pymmcore-plus/pymmcore-plus.

(58) Saini, A.; Messenger, H.; Kisley, L. Fluorophores “turned-on” by corrosion reactions can be detected at the single-molecule level. ACS Applied Materials & Interfaces 2020, 13, 2000–2006.

(59) Pinkard, H. et al. Pycro-Manager: open-source software for customized and reproducible microscope control. Nature Methods 2021, 18, 226–228.

(60) Hao, X.; Zhong, B.; Liao, Z.; Sun, L. Fast autofocus method for piezoelectric microscopy system for high interaction scenes. Microscopy Research and Technique 2023, 86, 773–780.

(61) Moukthika Autofocus using OpenCV: A Comparative Study of Focus Measures for Sharpness Assessment. OpenCV.org, 2025; https://opencv.org/autofocus-using-opencv-a-comparative-study-of-focus-measures-for-sharpness-assessme Accessed: 2026-02-16.

(62) Chalfoun, J.; Majurski, M.; Blattner, T.; Bhadriraju, K.; Keyrouz, W.; Bajcsy, P.; Brady, M. MIST: accurate and scalable microscopy image stitching tool with stage modeling and error minimization. Scientific Reports 2017, 7, 4988.

(63) The MathWorks, Inc. Image Processing Toolbox. version R2024a, The MathWorks, Inc.: Natick, Massachusetts, United States, 2024.

(64) MathWorks estgeotform2d: Estimate 2-D geometric transformation from matching points. https://www.mathworks.com/help/vision/ref/estgeotform2d.html, 2024; Accessed: 2026-02-18.

(65) Glira, P.; Weidinger, C.; Otepka-Schremmer, J.; Ressl, C.; Pfeifer, N.; Haberler-Weber, M. Nonrigid point cloud registration using piecewise tricubic polynomials as transformation model. Remote Sensing 2023, 15, 5348.

(66) Ma, J.; Kim, J. M.; Hoque, M. J.; Thompson, K. J.; Nam, S.; Cahill, D. G.; Miljkovic, N. Role of thin film adhesion on capillary peeling. Nano Letters 2021, 21, 9983–9989.

(67) Shaib, A. H. et al. One-step nanoscale expansion microscopy reveals individual protein shapes. Nature Biotechnology 2024, 43, 1539–1547.

(68) Mishra, S.; Scarano, F. J.; Calvert, P. Entrapment of Saccharomyces cerevisiae and 3T3 fibroblast cells into blue light cured hydrogels. Journal of Biomedical Materials Research Part A 2012, 100A, 2829–2838.

