## Supplementary material for "Tensile expansion microscopy applies mechanical force to super-resolve fixed and image live cellular samples": Supplimentary Information

### Methods

#### Image registration of macroscale fiducial markers

Expansion factors and the isotropy of expansion quantify the post expansion resolution and error in uniform expansion. The expansion factor is typically determined macroscopically, by comparing the size of the post expansion and pre expansion gel.[1] Alternatively, some ExM methods quantify the expansion factor microscopically by comparing the pre and post expansion sizes of reference structures like microtubules, clathrin coated pits or other nano structures that have well known size.[1,2] Local non uniform deformation in the expansion of the gel induce distortion of the image at a macroscopic and microscopic level. A level of anisotropy is inherent in the system due to the heterogeneity of the polymer matrix and is compounded by the application of non uniform stress to the system. Typical ExM protocols characterize this distortion by calculating the deformation vector field through a nonrigid image registration process.[1,3,4] From the vector field, a root mean square (RMS) error is quantified. Subsequent protocols have been established that using fiducial markers on the gel as rulers to monitor the microscopic expansion and local distortion by having a uniform distribution of the markers per area that are independent from the sample of interest.[3,4]

Here, we demonstrate our image registration analysis while using macroscopic fiducial markers to obtain the local deformation and quantify the local expansion factors on a macroscopic scale. A video of the expansion was collected on a Fujifilm XT3 with a Fujifilm XF 16-50mm f/2.8-4.8 R LMWR lens at 24 frames per second in grey scale. The gel was loaded onto the iris expansion device. Circular fiducial markers used were cut from black paper using a 1/16 inch hole punch. The markers were placed in roughly a grid shape for convenience. The iris expansion device was placed over a commercial stencil back light to improve contrast between the fiducial markers and the background. Since the fiducial markers are simply placed on top of the gel and not integrated into it, and since they are not elastic, the markers do not stretch with the gel and only serve to mark relative positions as the gel stretches in this case as seen in Fig. S3a,b.

Ideally the expansion can be mathematically expressed as a rigid similarity transform which includes translation, rotation and scaling. In reality, the expansion is not perfectly isotropic due to local non uniform deformations. The deviation from isotropy can be mathematically represented as a non-rigid transformation.[1 5] A post-expansion image can be registered to a pre-expansion image exactly by first doing a similarity transform, which gives us the global scaling parameter (i.e., global expansion factor), and then doing a non-rigid transform to obtain the deformation vector field which can quantify the error in expansion though a RMS error.

Fiducial markers are first identified and tracked to associate each marker to itself on different frames over expansion. We binarize the grey scale image based on a threshold calculated using Ostu's method.[6] Blob Analysis from MATLAB's computer vision toolbox[7] is used to identify the markers on MATLAB R2024a. Once the markers are detected on all frames of the video (Fig. S3a,b) , a nearest neighbor algorithm is used to track the markers through the expansion and generate trajectories. Then a triangulation (inbuilt Matlab function delaunayTriangulation) is applied to the grid of fiducial markers to obtain smaller regions to quantify expansion factors locally at 100 µm scales (Fig. S3c,d) and compare to the global expansion factor over 10 mm scales.

The expansion factor can be obtained in multiple ways at different length scales and compared to realize quantitatively the isotropy of the expansion. A global estimate of the expansion factor can be determined by comparing the size of the gel before and after expansion. Another way to estimate the global expansion factor would be to apply a similarity transform (including translation, rotation and scaling) of the post-expansion image to the pre-expansion image. This is done based on the fiducial markers identified through blob detection and particle tracking analysis. The similarity transform was estimated using an inbuilt MATLAB function estgeotform2d on MATLAB R2024a. The transform parameters extracted include a scaling factor which gives a calculated estimate of the global expansion factor. The obtained transform is then applied to the post expansion image (Fig. S4b) with respect to the pre-expansion image (Fig. S4a) to obtain the transformed image in Fig. S4c. The transformed final image and the before image are overlaid in Fig. S4d. The area and linear expansion factor can then be evaluated for smaller domains by comparing either the area or side lengths of the triangles, respectively, in the triangulation applied to the fiducial markers on the pre- and post-expansion images.

We obtain the deformation field corresponding to the non-rigid part of the deformation between the pre- (Fig. S4b) and post- (Fig. S4a) expansion images based on a triangulation based linear interpolation (Fig. S5a).[5] The deformation field represents the error or anisotropy of the equibiaxial expansion. To calculate the deformation field, we use the same triangulation on the fiducial markers generated before expansion as shown in Fig. S3 and maintain the triangulation through the expansion. Given a triangle with vertices v1v2v3, we know the position of the fiducial markers pre and post expansion with the similarity transform applied (represented in Fig. S5b by the magenta arrows). The difference between these points would be error in expansion. For any point p in the triangle, we will find the deformation based on a deformation field that is continuous and differentiable. This allows for a non-rigid but continuous movement of any point in the image internal to the fiducial markers. The vectors for the deformation field are calculated based on triangulation based linear interpolation. For a point p which is contained in triangle v1, v2, v3 there are some barycentric weights b1, b2, b3 respectively for each vertex. Multiplying the calculated displacements of the fiducial markers represented by v1, v2, v3 with the barycentric weights will give us the vector deformation at point p. That is p = v1b1 + v2b2 + v3b3. Repeating this for a grid of points gives us a representation of the deformation field as shown in Fig. S5a.

### Figures

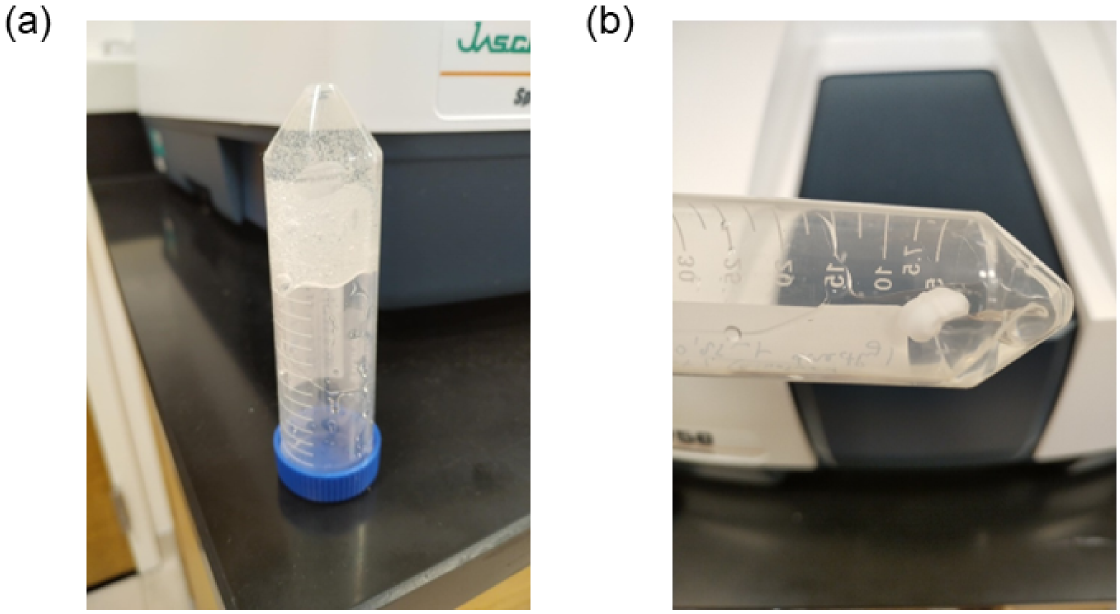


**Figure S1: Argon effect on AAm polymerization**: (a) The polymerization of AAm and the formation of PAAm covalently bonded network was achieved within 7 minutes of argon purging at room temperature to reduce oxygen quenching of the radical polymerization. (b) AAm was not polymerized after 15 hours at room temperature in the absence of argon purging

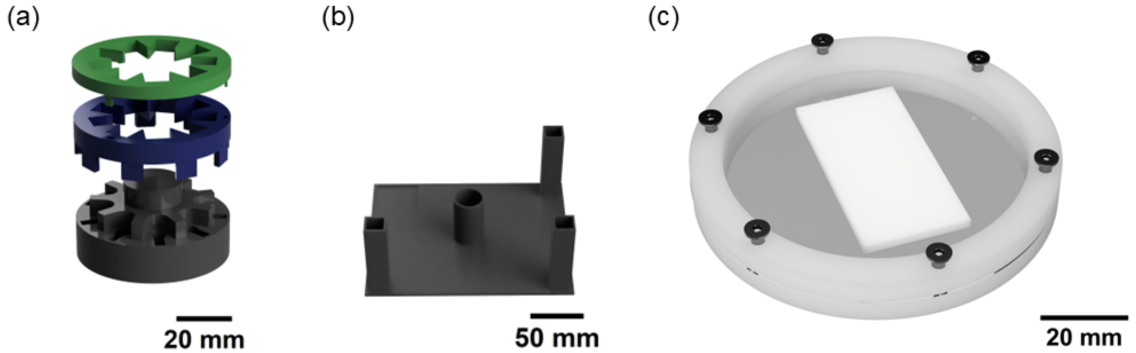


**Figure S2: Additional parts of the iris expansion device for sample preparation and preservation.** (a) Hydrogel mount (b) stand (c) drying ring. The gel mount and support stand were engineered to facilitate seamless sample loading onto the iris expansion device, prioritizing ergonomic handling and procedural repeatability. According to Fig. S2a, the bottom gel mount (black) has slots that fit the arms of the iris expansion device while the top parts (blue and green) were used to lock the grippers in place while loading the hydrogel sample. The hydrogel was placed on top of the bottom gel mount followed by the grippers then top parts (blue and green). Due to focal distance of the microscope's objective, the sample needs to be loaded below the arms which require the iris expansion device to be inverted on the stand (Fig. S2b) while loading the sample. The drying ring is designed to preserve the expanded hydrogel sample by preventing it from shrinking while drying at room temperature, as shown in (c), area of interest on the hydrogel sample is sandwiched between two glass coverslips. All parts are 3D printed with PLA material using Bambu Lab H2S 3D printer.

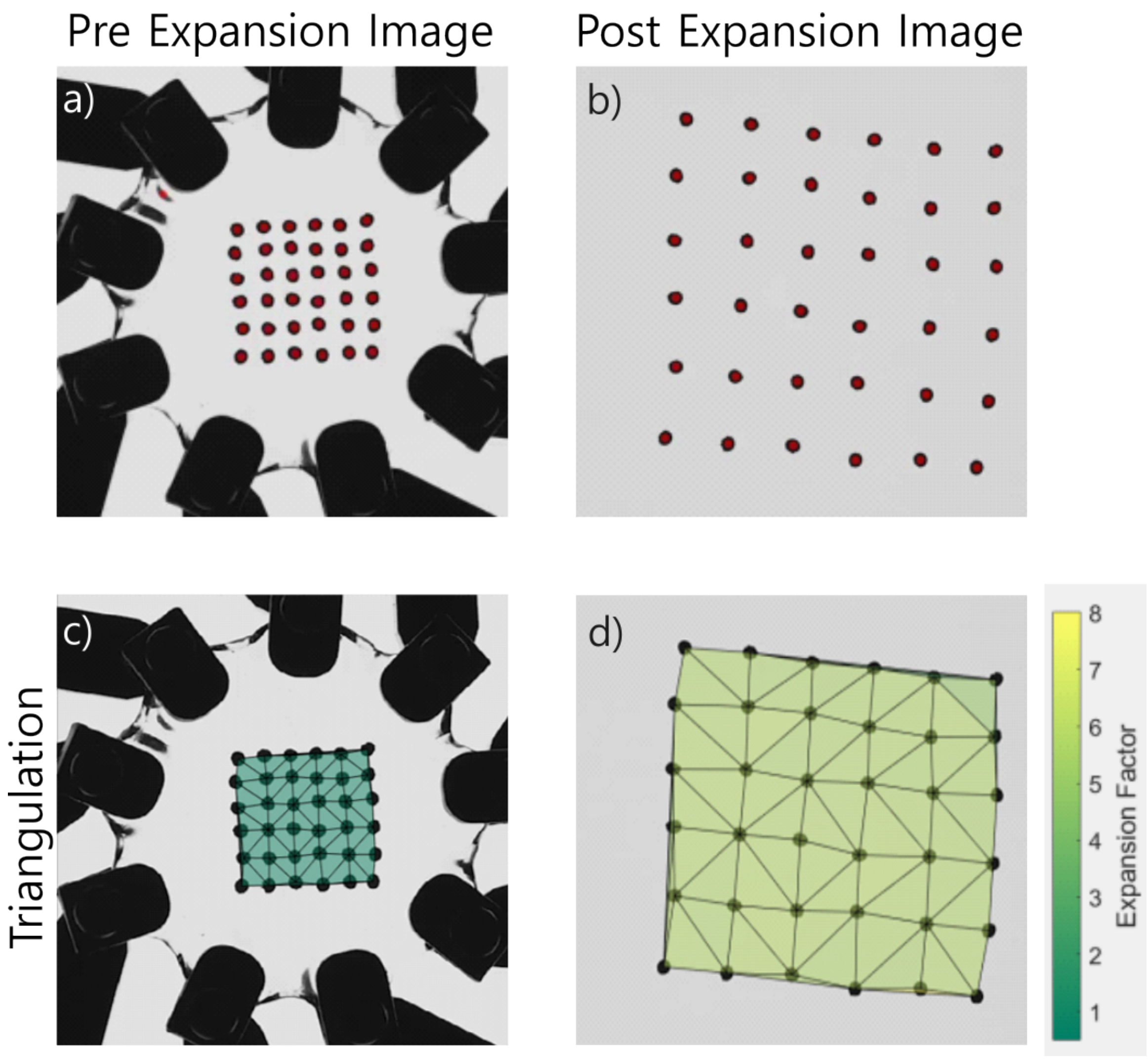


**Figure S3: Identification of fiducial markers and triangulation applied to the markers to obtain expansion factors for pre and post expansion images.** (a) Pre-expansion image with the centroid of the identified fiducial markers highlighted in red. (b) Post-expansion image with the identified fiducial markers highlighted in red. (c) Triangulation applied to the identified fiducial markers in the pre-expansion image with the local expansion factors represented by the fill color of the regions. Prior to expansion, all areas have a factor of 1. (d) Same triangulation applied to the fiducial markers on the post expansion image with the local expansion factors represented by the fill color of the region.

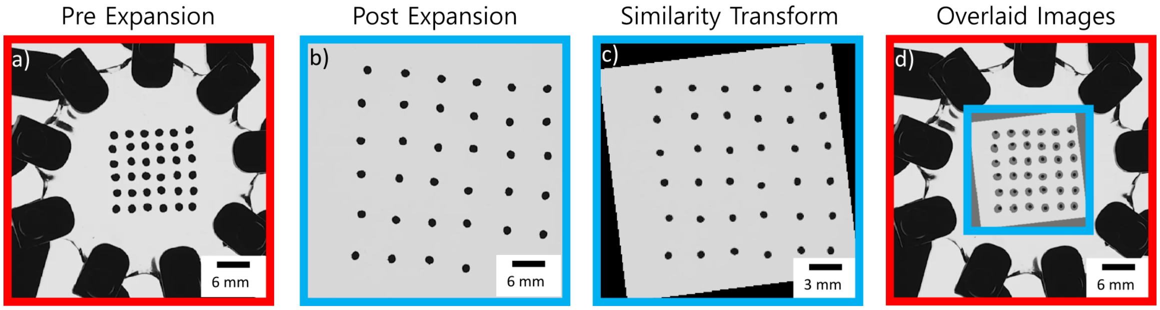


**Figure S4: Steps of image registration.** (a) Pre-expansion image of fiducial markers, (b) post-expansion image of fiducial markers, (c) similarity transform of the post-expansion image with respect to the pre-expansion image, (d) transformed post-expansion image overlaid with the pre-expansion image. Red and blue borders are used to indicate the same image and relative size/position when overlaid.

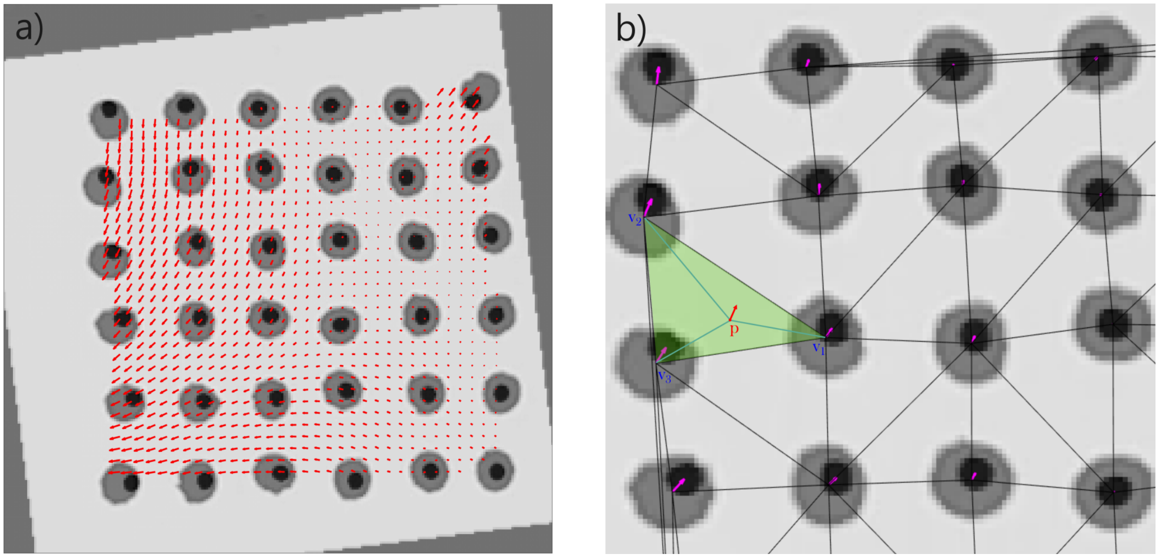


**Figure S5: Representation of the vector field deformation estimated through linear interpolation.** (a) Overlaid image with the deformation vector field represented by red arrows. (b) Portion of fiducial markers of the pre-expansion image and post expansion image with the transformation applied overlaid. Magenta arrows (going from pre-expansion image to the post expansion transformed image) represent the mismatch between the two sets of images. Smaller arrows represent smaller local errors in the equibiaxial stretch.

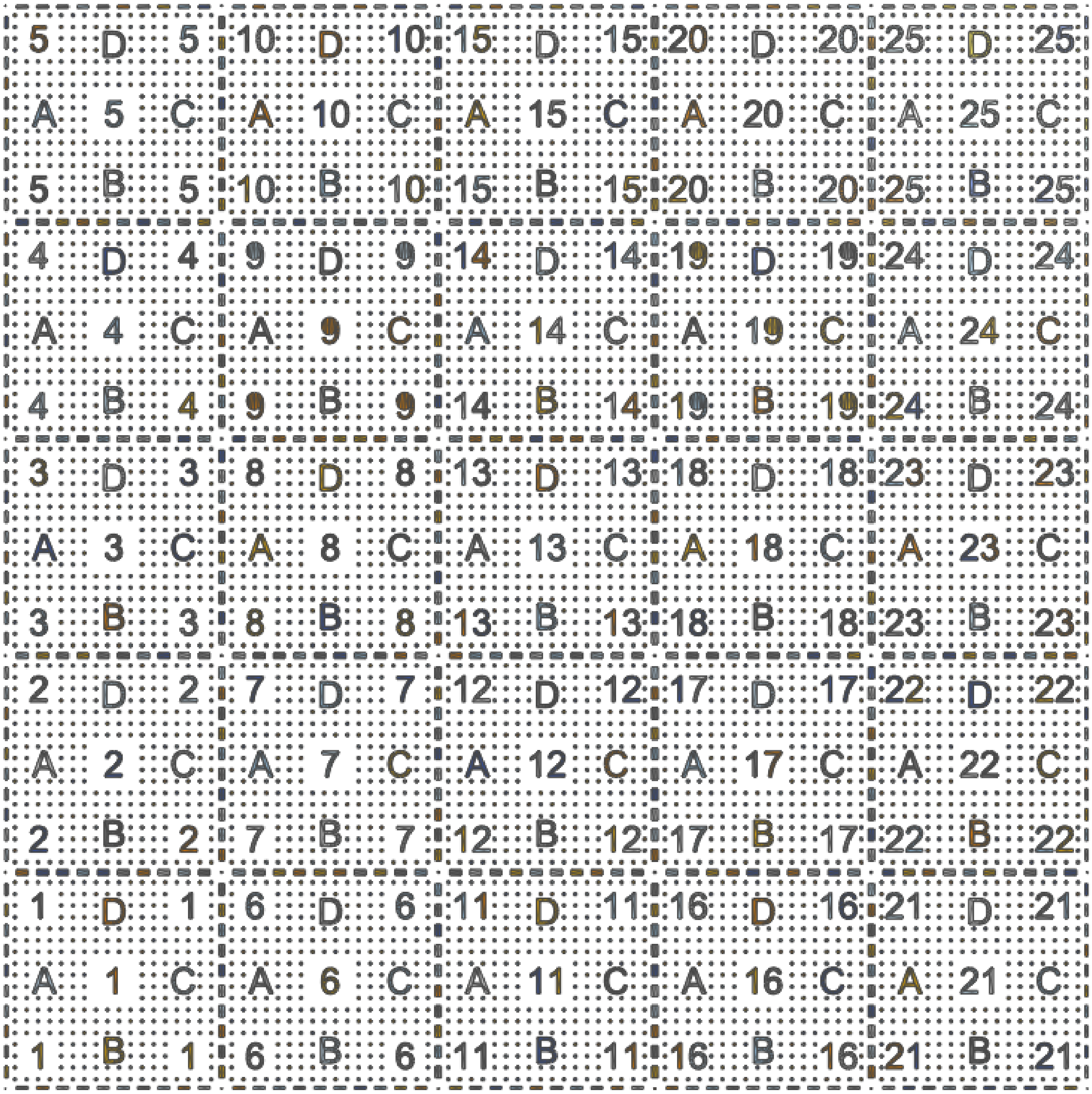


**Figure S6: The design of the fiducial markers using Computer-aided Software (CAD).**

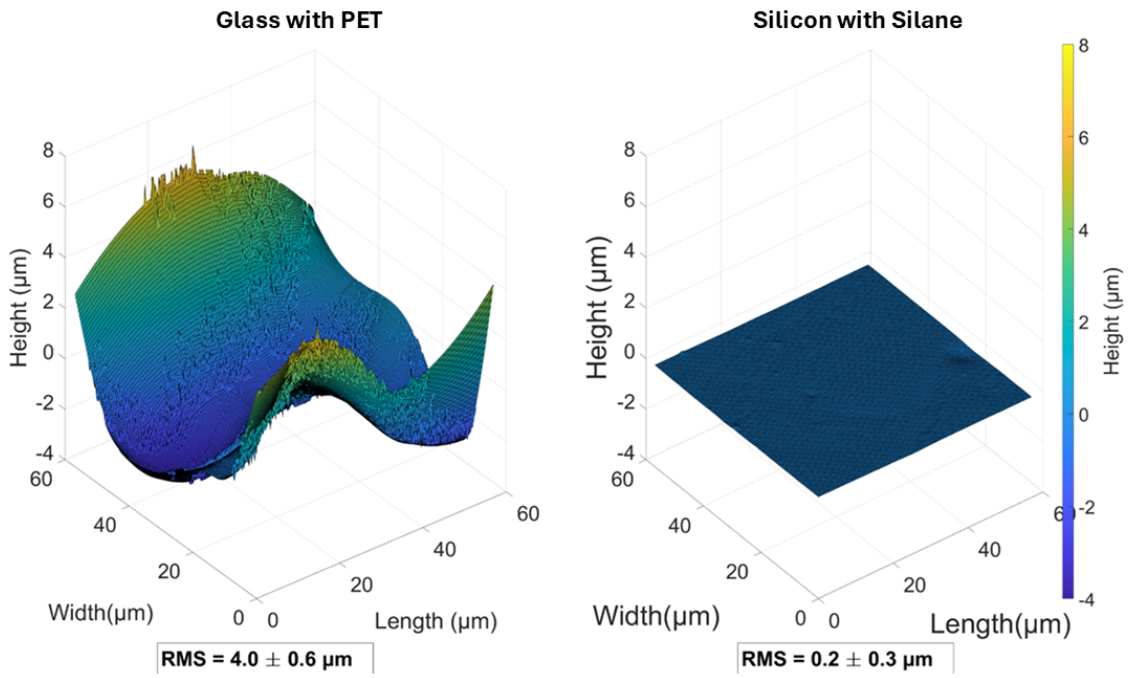


**Figure S7: Surface roughness analysis comparison between hydrogels polymerized on different substrates**. Hydrogels were polymerized on glass slabs covered by thin crystalline PET sheets and on silicon wafers coated with perfluorodecyltrichlorosilane. Both substrates are hydrophobic to minimize influence of substrate/hydrogel interaction on hydrogel surface roughness. Optical profilometry of the surfaces of these hydrogels show non-uniform surface for the hydrogels polymerized on glass with PET on the order of 4 $\mu$m, while the roughness of the hydrogels polymerized on the silanated wafers was measured to be in the order of 200 nm. We find that surface flatness is crucial for imaging to keep fluorescence features in the depth of field of the objective.

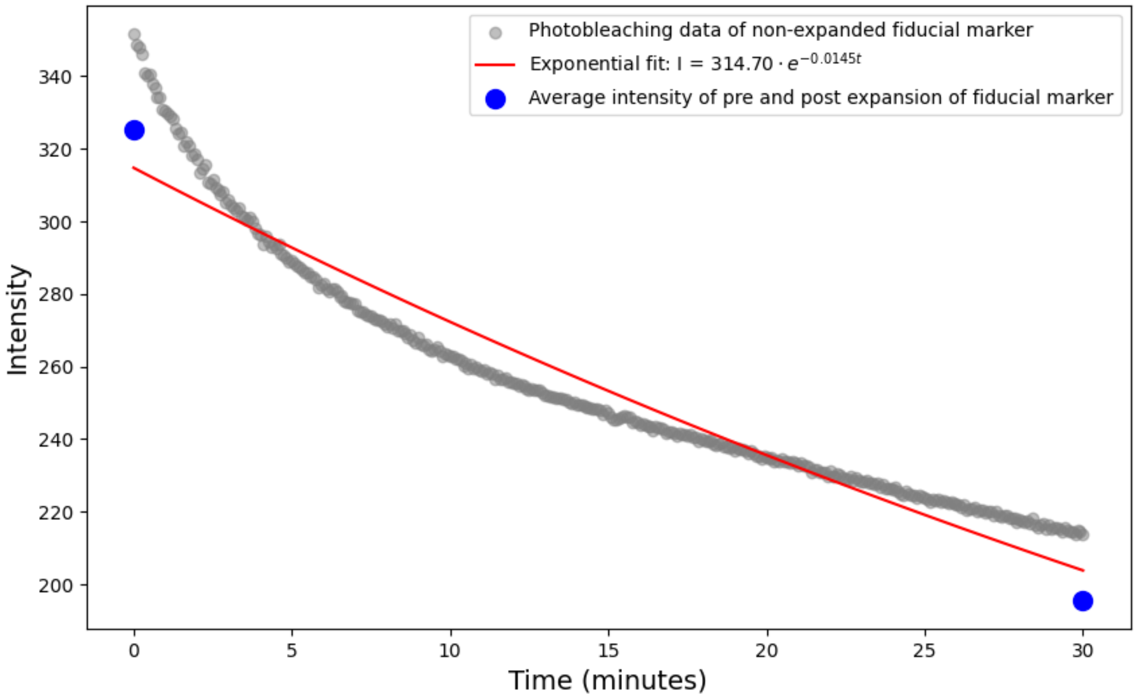


**Figure S8: Comparison between average intensity change of pre (1×) and post (3×) fiducial markers expansion on the hydrogel (16 wt.%) and non-expanded fiducial marker on hydrogel after a long-exposure to 633 nm laser (5 mW, 10 ms).** We observed lower intensity counts in pre- and post-expansion images. The result shows a decrease in average intensity of pre- and post-expansion which follows the photobleaching trend of the non-expanded fiducial marker. Thus, the result confirms the photobleaching of fiducial marker is due to longer exposure to laser rather than dye diffusion into hydrogel.

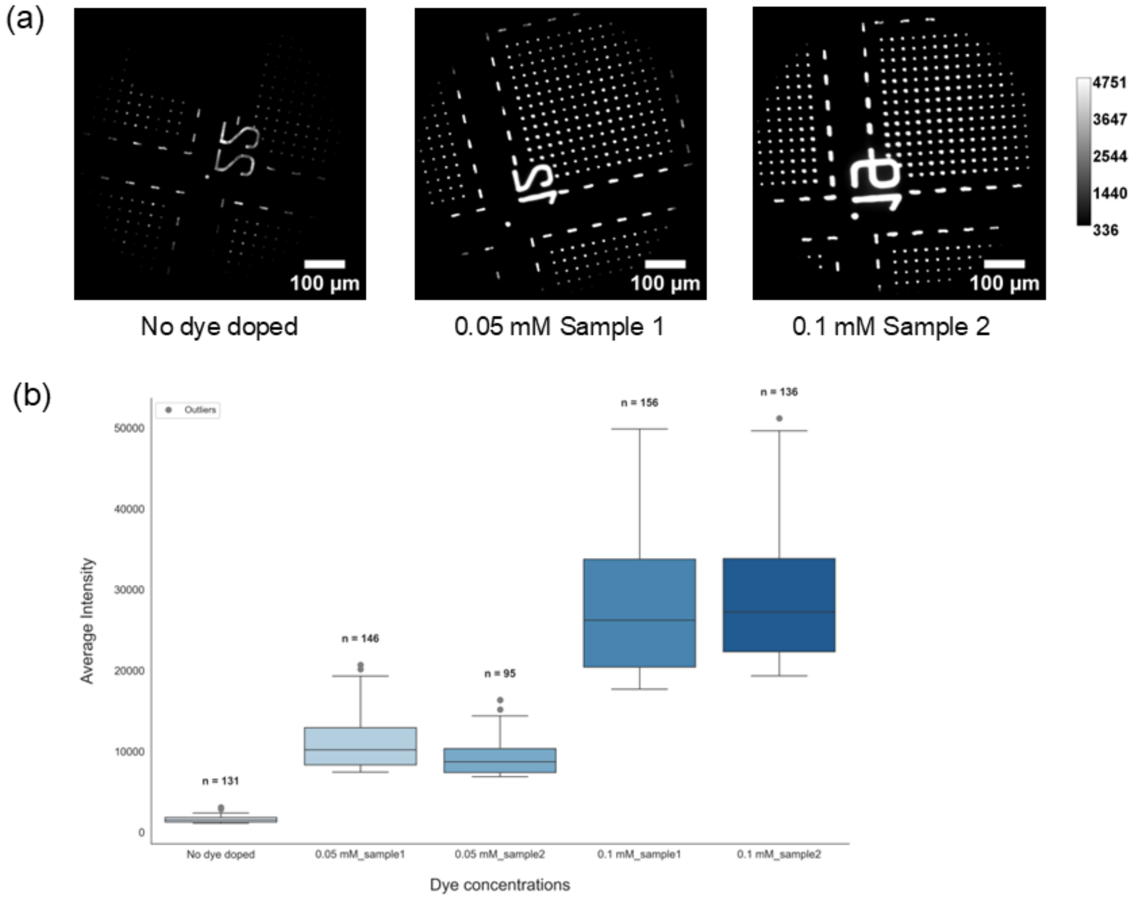


**Figure S9: Intensity measurement of Nanoscribe IP-S doped with different concentrations of rhodamine 6G dye demonstrates versatility of two-photon lithography fiducial markers for analyte doping.** (a) Example images of the fiducial markers doped with different rhodamine 6G dye concentrations. All images are displayed at the same intensity scale for comparison. Fluorescence intensity increases at higher dye concentration. (b) A box plot of the average intensity of each doping conditions. We observe a 19-fold increase in average intensity count between non-doped and 0.1 mM doped sample while there is approximately 3-fold increase between 0.05 mM and 0.1 mM doped samples.

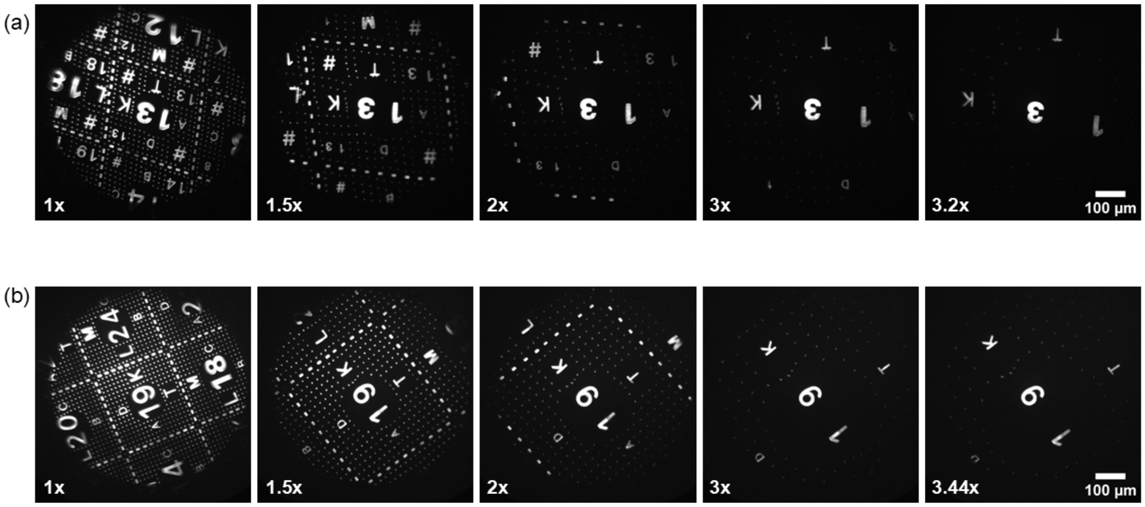


**Figure S10: Real time observation of Atto633-doped Nanoscribe IP-Visio resin during expansion.** Nanoscribe IP-Visio fiducial markers doped with 10 mM Atto633 is printed on silicon wafer substrate before polymerizing hydrogel (16 wt.%) as described in the methods. Real time image acquisition was performed with a 633 nm laser at 2.5 mW and 50 ms exposure time. Dynamic expansion using the iris expansion device using a step size of 0.04×, speed of 0.01 mm/sec, and additional 10 seconds dwell time after each step allowed the hydrogel to reach equilibrium. The features of interest (numbers 9 and 3, respectively for (a) and (b)) are focused and tracked to kept in the center of the field of view during expansion for visualizing real-time changes and performing expansion analysis. The images shown undergo constant scaling using Autoscale on ImageJ software. We notice photobleaching of the fiducial markers at higher expansion due to the continued, long exposure to the laser during the continuous imaging. Videos of real time expansion are attached as "Video_S1a.gif, Video_S1b.gif" found in our github linked below.

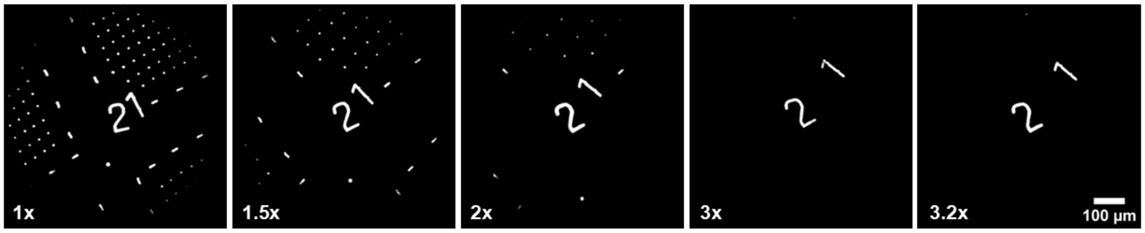


**Figure S11: Real time observation of alternative Nanoscribe IP-S resin that is observable with 561 nm excitation during expansion due to autofluorescence.** Here we demonstrate an alternative Nanoscribe IP-S on a silicon wafer substrate before polymerizing hydrogel can also serve as fiducial markers. The IP-S is transferred to the hydrogel (16 wt.%) in the same manner as the IP-Visio as described in the methods. Real time image acquisition with an interested fiducial marker (number 2) being focused and tracked in the center throughout the expansion. A 561 nm laser at 5 mW and 10 ms exposure time was used to acquire the images. Dynamic expansion of the iris expansion device used a step size of 0.05x, speed of 1 mm/sec, and additional 10 second dwell time after each step allowed the hydrogel to reach equilibrium. The images shown undergo constant scaling using Autoscale on ImageJ software. We also noticed photobleaching on the fiducial markers with higher expansion due to the longer exposure. In contrast to the IP-Visio which required doping with Atto633 to observe (Fig. S10), the IP-S resin has autofluorescence at 488, 561, and 633 nm excitations. Autofluorescence can have a benefit of not requiring doping of fluorophores but with the drawback of potential overlapping emission with dye labels or fluorescent proteins within cellular samples. Videos of real time expansion are attached as "Video_S2.gif" found in our github linked below.

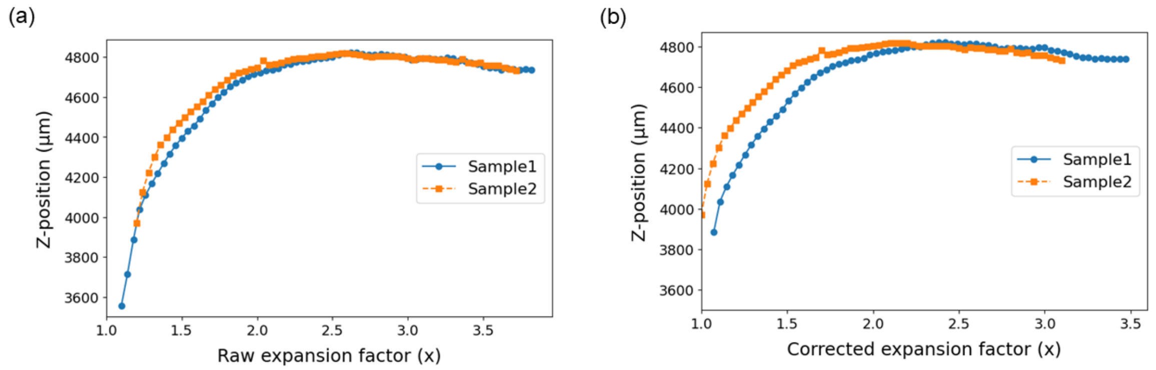


**Figure S12: The hydrogel thins in axial dimension during lateral expansion.** Plotting the objective focal position (z-position) based on the microscope metadata vs. expansion factor confirms the reduction in thickness of hydrogel (16 wt.%) as expansion factor increases. (a) z-position vs raw expansion factor and (b) corrected expansion factor due to the difference in the initial expansion setting of the two samples. The "raw" expansion factor setting, which is the value displayed on the iris device electronics can differ due to sample tautness resulting from sample loading step. The thickness of fully stretched hydrogel is determined from initial hydrogel thickness (1.5 mm) minus the change in focus position (z-position) between pre-stretch and post-stretch hydrogel. The final hydrogel thickness is approximately 319 μm, calculated from Sample 1 in Fig. S12a due to data completeness which resembles ideal stretching condition. The changes in hydrogel thickness motivate the need for the autofocus software (as described in Methods) due to limitation of typical hardware autofocus which require the sample to have consistent focal position.

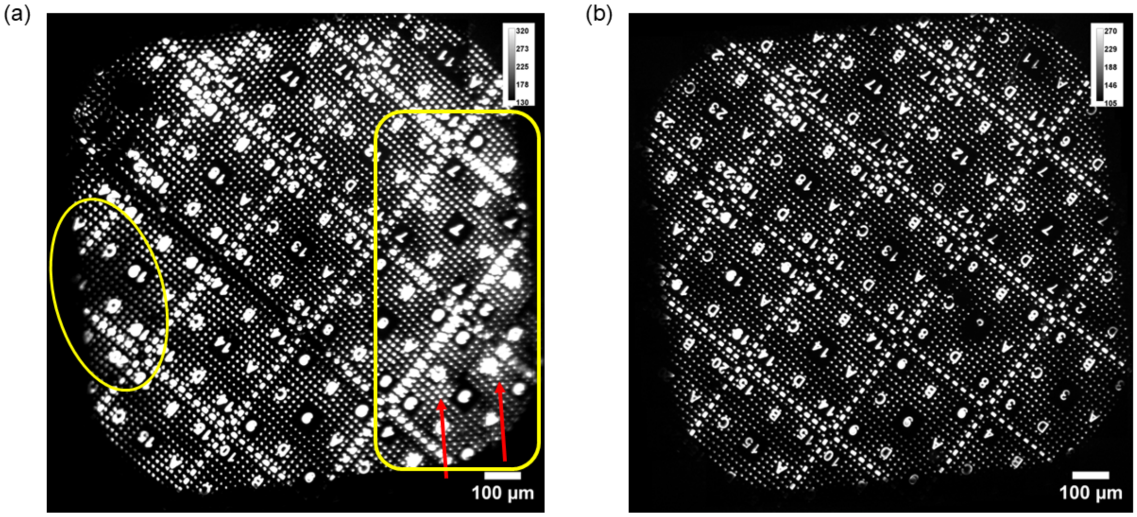


**Figure S13: Necessity of autofocus algorithm due to differences in focal plane.** (a) Failure of multiple field-of-view images stitching of fiducial markers without autofocus due to difference in focal plane across the sample. Red arrows illustrate visible stitching lines resulting from image registration's inaccuracy due to out-of-focus area of the fiducial markers (yellow boxes). Thus, autofocus is required after for each frame during the multiple field-of-view acquisitions. (b) shows successful stitching result of the fiducial markers after incorporating the autofocus software.

(a)

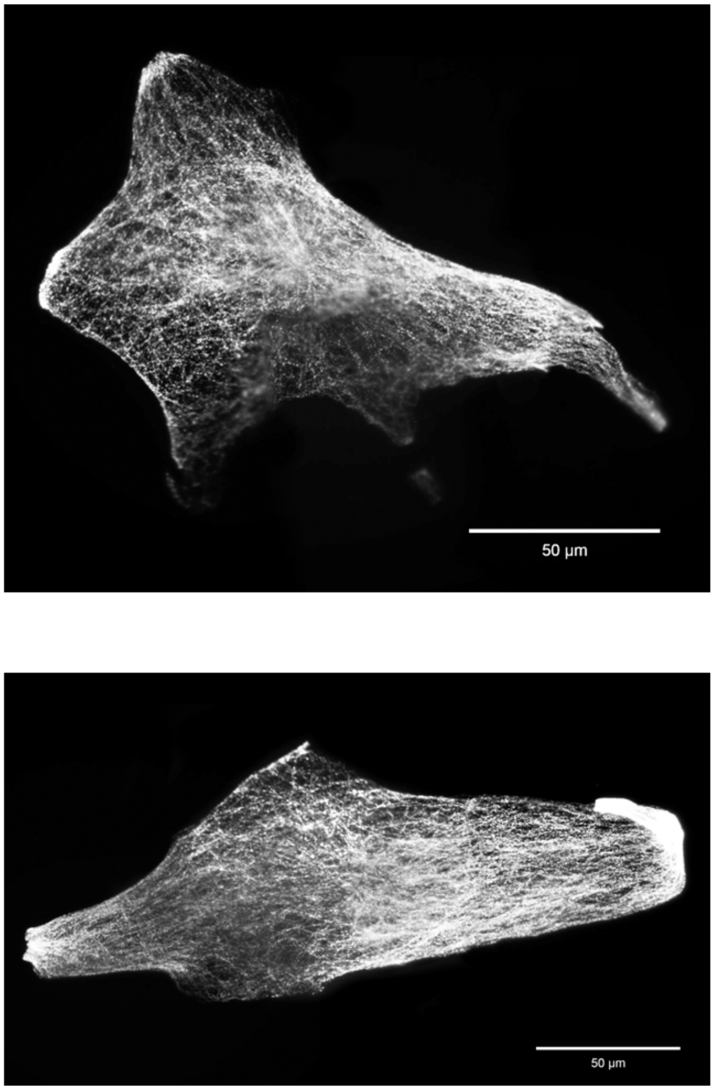


(b)



**Figure S14: Expanded fixed 3T3 cells observed with a 100x objective.** Figure S14a,b demonstrates two fully expanded fixed 3T3 cells images acquire using 100x objective (1.49 NA) with a 488 nm laser at 3.54 mW and 100 ms exposure time. A total of 20 frames were taken and summed together to increase signal counts. Both samples were prepared and stained as described in the methods. After the samples were fully expanded on the iris expansion device, a drying ring (as shown in Fig. S2c) and glass coverslips were used to preserve the sample. The resulting data for a single expanded cell were taken from multiple fields of views (100 μm x 100 μm) and were stitched together using Photoshop software. Different areas of the cells also spanned different focal planes in the axial dimension leading to some blur in (a).

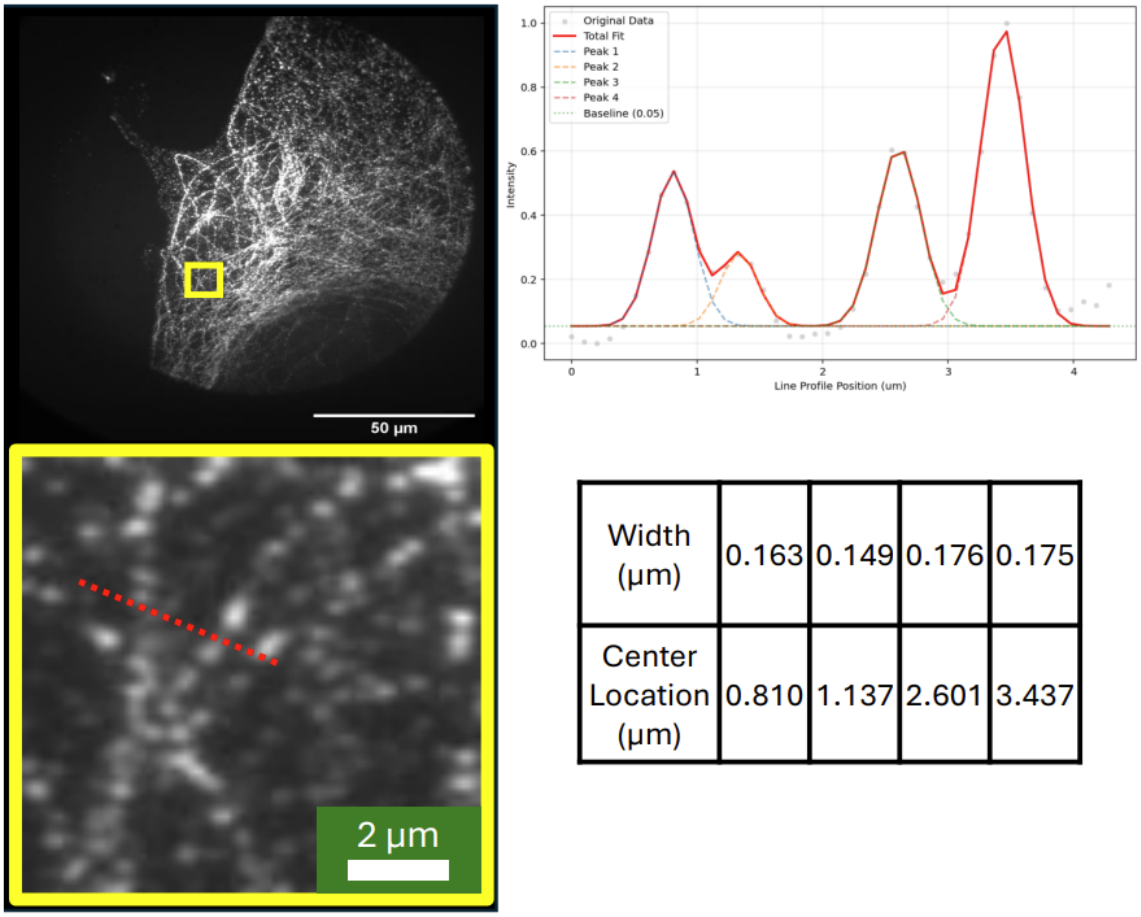


| Width (µm) | 0.163 | 0.149 | 0.176 | 0.175 |
|---|---|---|---|---|
| Center Location (µm) | 0.810 | 1.137 | 2.601 | 3.437 |

**Figure S15: Fixed 3T3 cell microtubles analysis demonstrates super-resolution cell image.** Fixed 3T3 cell was expanded and dried as mentioned in the Method. Image in the blue box is cropped from its full image on the left obtained from 100x objective to demonstrate line section (orange) on 3T3 cell's microtubulars. From the line section, Gaussian-fitted plot of intensity reveal our capability of achieving super-resolution image through tensile expansion microscopy (TExM).

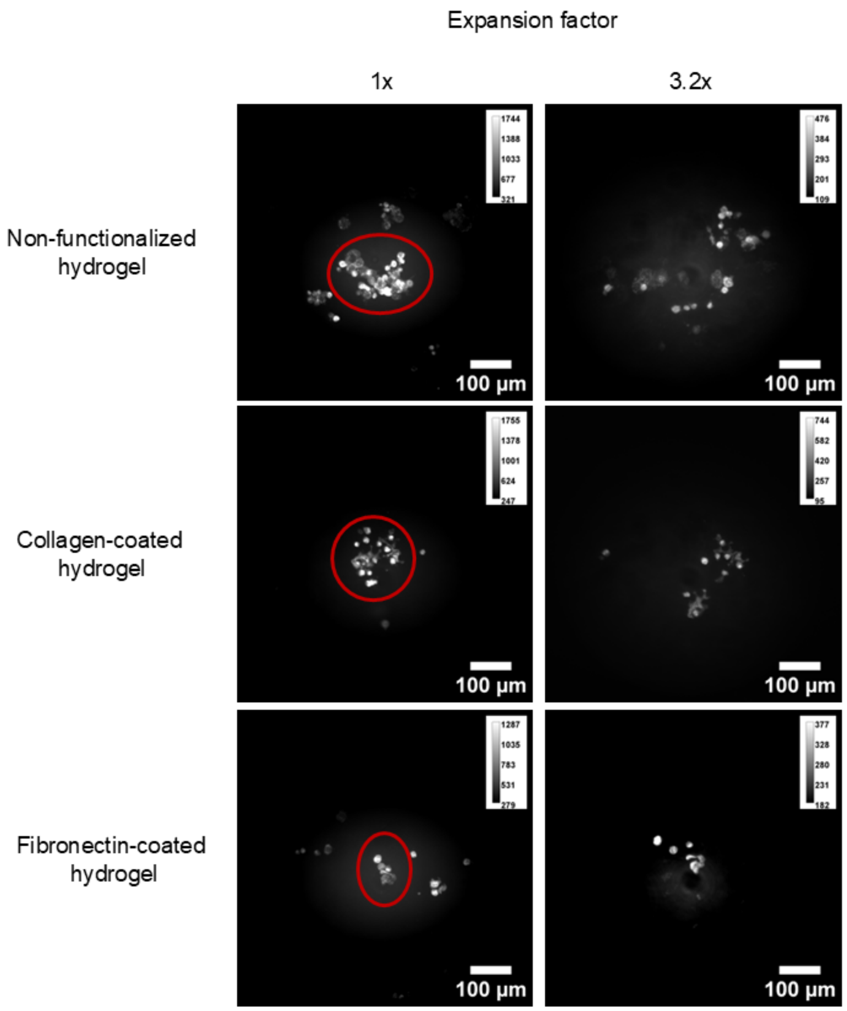


**Figure S16: Observation of the effect of hydrogel surface functionalization live HeLa cell seeding.** Hydrogels (14 wt.%) were made and sterilized according to protocol mention in the methods then coated with different cell adhesion proteins: collagen and fibronectin, prior to HeLa cell seeding. For collagen-coated hydrogels, 200 µg/mL of collagen was obtained from diluting stock collagen I from rat tail (Prod #: 354249, CAS #: 9007-34-5, Corning) in DPBS and NaOH was added dropwise until a pH of approximately 7.4 was reached. 200 µg/mL collagen was dropped on the hydrogel surface then incubated for 1 hour at 37°C and 5% $CO_2$. To coat fibronectin on hydrogel's surface, 2 mM of Sulfo-SANPAH (Prod#: 22589, CAS #: 102568-43-4, Thermo Fisher Scientific) was dropped and activated using UVA (365 nm) for 10 minutes prior to addition of 200 µg/mL fibronectin (Prod#: 33016015, Thermo Fisher Scientific) then incubated for 1 hour at room temperature. All hydrogels were seeded with the same density (250,000 cells in 200 $\mu L$ of cell media) of HeLa cells and incubated at 37°C for 1 day. The figure shows the results of cell adhesion on each adhesion proteins compared to non-functionalized hydrogel. We observed higher confluency (more adhesion) of HeLa cells on non-functionalized hydrogel and there were no significant difference in cells' morphology across the samples. Comparing between pre- and post-expansion of functionalized and non-functionalized hydrogels, the cells adhered to non-functionalized hydrogel are more stretched apart. Therefore, we proceeded with non-functionalized hydrogel for live HeLa cells expansion experiment.

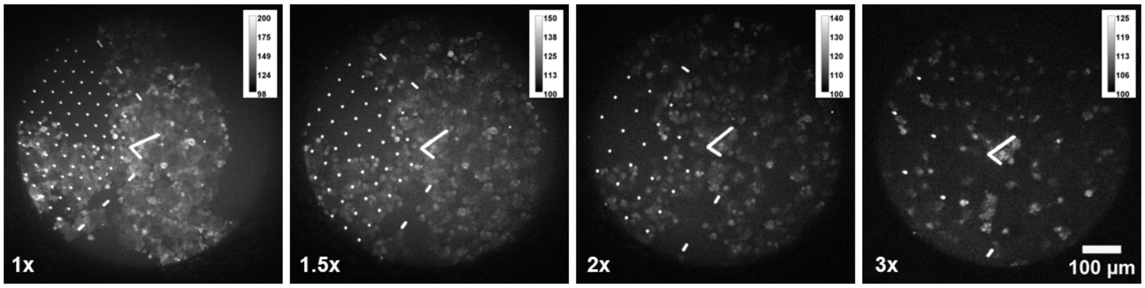


**Figure S17: Real time observation of HeLa cells separating during expansion with Nanoscribe IP-S resin fiducial markers present.** Real-time image acquisition of Nanoscribe IP-S fiducial markers and live HeLa cells seeded on hydrogel (14 wt.%) being tracked throughout the expansion. Image acquisition used 488 nm excitation at 5 mW and 20 ms exposure time. The iris expansion device used a step size of 0.05×, speed of 0.01 mm/sec and additional 10 seconds dwell time after each stretch to allow the hydrogel to equilibrate. We found that expansion rate affects the adhesion of the cells to the hydrogel substrate, as faster stretching speed led to cell detachment from the hydrogel. Comparing the pre (1×) and post-expansion (3×) images, photobleaching occurred at a higher magnitude for the cells compared to the fiducial markers which reduces their visibility at higher expansion. In addition, we observed several changes in cells behavior. For instance, cell clusters move apart from each other and some cells burst open. Videos of real time live HeLa cells expansion are attached as "LiveHeLa_video_S3.gif" found in our github linked below.

### Tables

Table S1: Iris Expansion Measurements

| **Iris Expansion** | **Across** | **Expanded Ratio** | **Down** | **Expanded Ratio** |
|---|---|---|---|---|
| 1.0x | 27 µm | 1 | 49 µm | 1 |
| 1.7x | 39 µm | 1.4 | 65 µm | 1.3 |
| 2.1x | 43 µm | 1.6 | 77 µm | 1.6 |
| 2.5x | 46 µm | 1.7 | 101 µm | 2.1 |
| 3.2x | 49 µm | 1.8 | 120 µm | 2.4 |

Table S2: Description of sample conditions

| **Figure** | **Fiducial marker** | **Substrate** | **Hydrogel** | **Uplifting** | **Calcium soak** | **Digestion** |
|---|---|---|---|---|---|---|
| Figure 1 | 2 mm hole punch cutouts | Crystalline PET sheet | Hydrogel (14 wt.%) | NA | 7, 8 minutes | NA |
| Figure 2 | 10 mM Atto-633 IP-Visio | Silicon wafer | Hydrogel (16 wt.%) | NA | 6, 6 minutes | NA |
| Figure 3 | 10 mM Atto-633 IP-Visio | ITO-coated 1.5 glass | Hydrogel (16 wt.%) | Uplifting buffer and physical peeling | 6, 6 minutes | Proteinase k 0.3 mg/mL |
| Figure 4 | 10 mM Atto-633 IP-Visio | Silicon wafer | Hydrogel (16 wt.%) | NA | 6, 6 minutes | NA |

**Analysis script, CAD files and gifs referenced in the SI can be found at the following GitHub repository**: https://github.com/KisleyLabAtCWRU/TExM

### References


(1) Chen, F.; Tillberg, P. W.; Boyden, E. S. Expansion microscopy. *Science* **2015**, *347*, 543–548.

(2) Damstra, H. G.; Mohar, B.; Eddison, M.; Akhmanova, A.; Kapitein, L. C.; Tillberg, P. W. Visualizing cellular and tissue ultrastructure using Ten-fold Robust Expansion Microscopy (TREx). *Elife* **2022**, *11*, e73775.

(3) Damstra, H. G.; Passmore, J. B.; Serweta, A. K.; Koutlas, I.; Burute, M.; Meye, F. J.; Akhmanova, A.; Kapitein, L. C. GelMap: intrinsic calibration and deformation mapping for expansion microscopy. *Nature Methods* **2023**, *20*, 1573–1580.

(4) Nakamoto, M. L.; Forró, C.; Zhang, W.; Tsai, C.-T.; Cui, B. Expansion microscopy for imaging the cell–material interface. *ACS Nano* **2022**, *16*, 7559–7571.

(5) Glira, P.; Weidinger, C.; Otepka-Schremmer, J.; Ressl, C.; Pfeifer, N.; Haberler-Weber, M. Nonrigid point cloud registration using piecewise tricubic polynomials as transformation model. *Remote Sensing* **2023**, *15*, 5348.

(6) Otsu, N. A Threshold Selection Method from Gray-Level Histograms. *IEEE Transactions on Systems, Man, and Cybernetics* **1979**, *9*, 62–66.

(7) Inc., T. M. Computer Vision Toolbox: 24.1 (R2024a). 2022; `https://www.mathworks.com`.
